# The role of icIL-1RA type 1 in oral keratinocyte senescence and the development of the senescence associated secretory phenotype

**DOI:** 10.1101/2020.07.06.189019

**Authors:** Sven E Niklander, Hannah L Crane, Lav Darda, Daniel W Lambert, Keith D Hunter

## Abstract

There is compelling evidence that senescent cells, through the senescence-associated secretory phenotype (SASP), can promote malignant transformation and invasion. IL-1 is a key mediator of this cytokine network, but the control of its activity in the senescence program has not been elucidated. IL-1 signalling is regulated by IL-1RA, which has four variants. Here, we show that expression of intracellular IL-1RA type 1 (icIL-1RA1), which competitively inhibits binding of IL-1 to its receptor, is progressively lost during oral carcinogenesis *ex vivo* and that the pattern of expression is associated with keratinocyte replicative fate *in vitro*. We demonstrate icIL-1RA1 is an important regulator of the SASP in mortal cells, as CRISPR-CAS9 mediated icIL-1RA1 knockdown in normal and mortal dysplastic oral keratinocytes is followed by increased IL-6 and IL-8 secretion, and rapid senescence following release from ROCK inhibition. Thus, we suggest that downregulation of icIL-1RA1 in early stages of the carcinogenesis process can enable the development of a premature and de-regulated SASP, creating a pro-inflammatory state in which cancer is more likely to arise.

## INTRODUCTION

The IL-1 receptor antagonist (IL-1RA) is an important regulator of IL-1 signaling in health and disease. IL-1RA acts as the main IL-1 inhibitor and has no agonist actions (Kurzrock et al., 2019), as it is a competitive antagonist of the IL-1 agonist receptor (IL-1R1), preventing IL-1 binding and the activation of downstream pathways (Hannum et al., 1990). IL-1RA is also able to inhibit the NF-κB and p38 MAPK transduction pathways in a non-IL-1R1 dependent manner (Garat and Arend, 2003). IL-1RA is coded by the *IL1RN* gene located at 2q14 (Eisenberg et al., 1990) and encodes four IL-1RA protein variants, one secreted (sIL-1RA) and three intracellular forms (icIL-1RA1-3), by alternate splicing (Haskill et al., 1991). sIL-1RA is a 17kDa protein secreted by monocytes and neutrophils (Malyak et al., 1998a). The intracellular variants lack a leader peptide thus cannot be secreted (Haskill et al., 1991). icIL-1RA1 is the main intracellular variant with a molecular weight of 18kDa and is mainly expressed in keratinocytes and other epithelial cells, monocytes, tissue macrophages, fibroblasts and endothelial cells. icIL-1RA2 protein has never been found in human cells (Arend, 2002). icIL-1RA3 has been found in monocytes, neutrophils and macrophages (Malyak et al., 1998b), but has 5-fold less affinity to IL-1R1 than the other forms and has not been reported in keratinocytes (Malyak et al., 1998a, Malyak et al., 1998b, Weissbach et al., 1998).

In senescent cells, IL-1 signalling is closely associated with the development of the senescence associated secretory phenotype (SASP)(Lau et al., 2019), a secretory state characterized by the production of more than 40 factors involved in intercellular signalling (Davalos et al., 2010). This includes a number of pro-inflammatory cytokines, including IL-6 and IL-8, molecules known to induce EMT and invasion of pre-malignant cells (Malaquin et al., 2013, Coppe et al., 2008, Ortiz-Montero et al., 2017, Lau et al., 2019). Thus senescence, whilst being a potent suppressor of malignancy in mutated premalignant cells (Kang et al., Nature 2011) is also considered to participate in the tumour promoting neoplastic environment (Campisi, 2005), as senescent cells are found in pre-malignant lesions *in vivo* (Guo et al., 2019), including those of the oral cavity (Natarajan et al., 2003). IL-1 signalling is important for the development of a microenvironment able to promote cancer development (Orjalo et al., 2009, Lau et al., 2019), as it regulates IL-6 and IL-8 *via* NF-κB activation (Wolf et al., 2001, Orjalo et al., 2009, Rovillain et al., 2011). Reduction in the levels of its natural antagonist, IL-1RA, has been related to tumour growth, invasion and metastasis (La et al., 2001, Lewis et al., 2006) and could allow a deregulation of the SASP (Acosta et al., 2013).

A number of transcriptomic studies have independently found *IL-1RN* to be downregulated in mucosal head and neck cancers (HNC) when compared to matched normal oral mucosa (Alevizos et al., 2001, Choi and Chen, 2005, Cromer et al., 2004, Leethanakul et al., 2000, Schmalbach et al., 2004, Whipple et al., 2004, Lallemant et al., 2009, Koike et al., 2005). Recently, Shiiba et al. (2015), reported total IL-1RA protein expression was progressively downregulated through oral dysplasia (OD) to oral squamous cell carcinoma (OSCC) (Shiiba et al., 2015), but its consequences are poorly understood. HNC comprise the 8th most common cancer worldwide and causes significant morbidity and mortality, with a 5-year survival rate of ≈ 50% (Marur and Forastiere, 2016). Oral cancer arises in many cases from oral potentially malignant disorders (OPMDs), which can present clinically as leukoplakias and erythroplakias (Villa et al., 2019). No treatment has been proven to successfully reduce malignant transformation of OPMDs (Lodi et al., 2016). Thus, it is important to identify the mechanisms involved in oral malignant transformation that could be pharmacologically targeted.

As IL-1RA downregulation is progressively observed through OD to OSCC, we have used this as a model to examine the contribution of IL-1RA to the malignant transformation process and to the development of a pro-tumourigenic SASP (which has not been characterized in oral keratinocytes), in order to understand how its downregulation contributes to the transformation towards HNC.

## RESULTS

### Intracellular IL-1RA expression is lost during carcinogenesis of oral keratinocytes

To assess *IL1RN* expression during oral carcinogenesis, *IL1RN* transcript levels were analysed in a panel of mortal normal oral keratinocytes (NOKs; NOK805, NOK829), immortalized normal oral keratinocytes (iNOKs: FNB6, OKF4 and OKF6), mortal and immortal dysplastic oral keratinocytes (OD: D6, D25 and D19, D20 respectively) and malignant oral keratinocytes (OSCC: B16 and B22). We observed a progressive decrease in *IL1RN* transcript levels in OD and OSCC cell lines compared to NOKs (Fig 1A and EV1A for individual cell expression which includes iNOKs). Total IL-1RA (tIL-1RA) protein levels were concordant with the qPCR data, showing a decrease in tIL-1RA protein in OD and OSCC cell lines compared to NOKs (Fig 1B). The decrease in tIL-1RA protein levels in OD and OSCC cells can be attributed to a decrease in mRNA levels of the intracellular variant (*icIL-1RN)* (Fig 1C), as no significant differences in expression of the secreted form (*sIL-1RN)* were observed (Fig 1D). The dominant isoform expressed in oral keratinocytes is *icIL-1RN* transcript variant 3 which encodes icIL-1RA type I (icIL-1RA1) (Fig EV1B). None of the immortal OD and OSCC cells showed any tIL-1RA expression (D19, D20 and B16 and B22, respectively), whereas tIL-1RA was readily detected in FNB6 (Fig 1B and E), in keeping with the transcript abundance. tIL-1RA was located diffusely across the cytoplasm and is present in the nucleus of FNB6 (Fig 1F). Immunohistochemical assessment of tIL-1RA expression in a panel of five normal oral mucosa, nine OD and ten OSCC patient tissue samples showed a reduction in expression in agreement with our *in vitro* data, with robust expression of tIL-1RA in normal oral epithelium and progressive loss of expression in OD and OSCC (Fig 1G and H).

**Figure 1:**
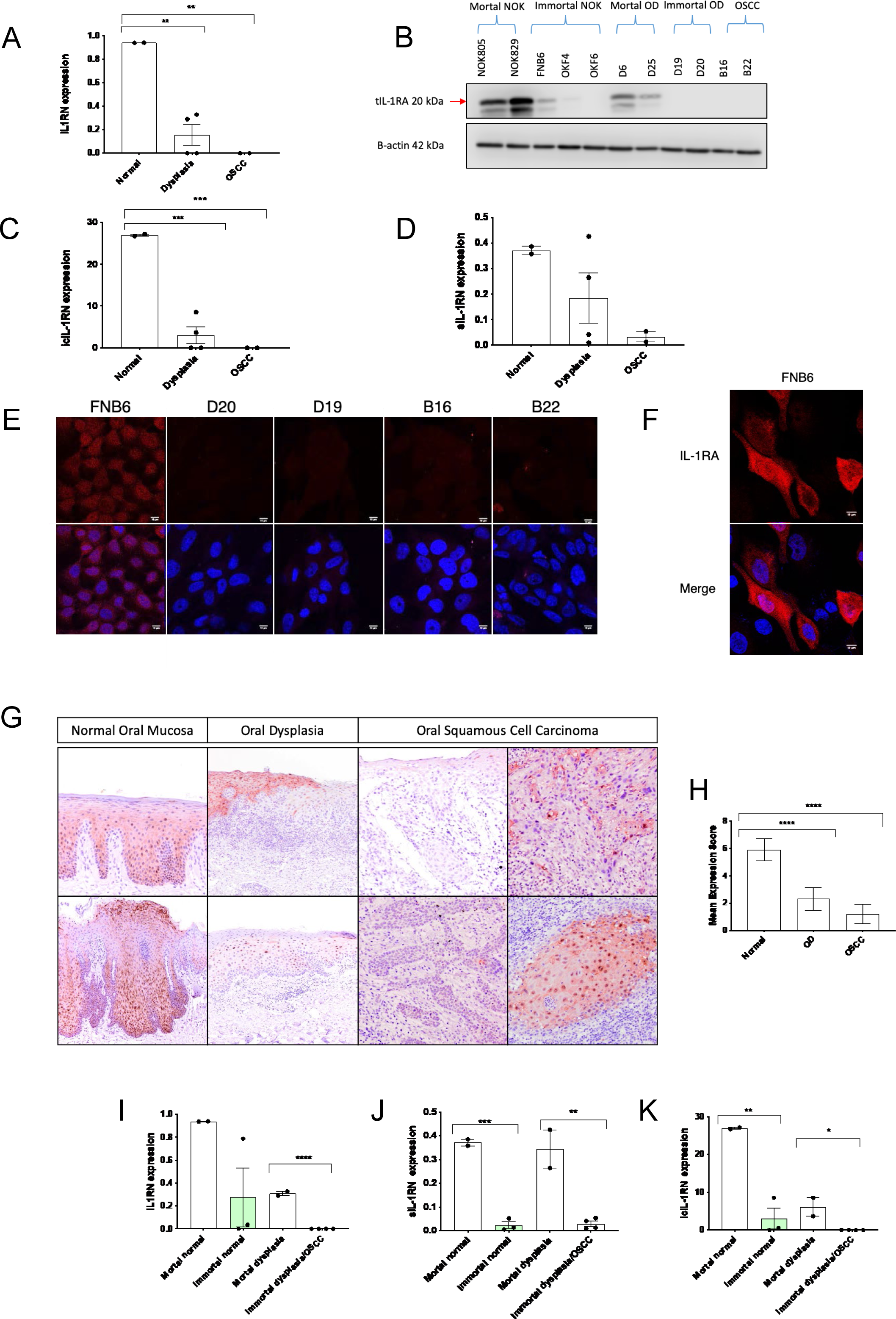
IL-1RA is downregulated during oral carcinogenesis *in vitro* and *in vivo*. A, C, D: Data obtained from two primary normal oral keratinocytes (NOK) cell cultures (NOK805, NOK829), four dysplastic cell cultures (D6, D25, D19, D20), and the two oral cancer cell lines (B16, B22) were clustered together into three groups: normal, dysplasia and oral squamous cell carcinoma (OSCC) respectively. Graphs showing IL1RN (A), icIL-1RN (C) and sIL-1RN (D) mRNA expression express as fold change. Data are shown as mean ± SEM (N = 3 independent experiments, n = 3 technical replicates). One-way ANOVA with multiple comparisons was used to calculate the exact P value. B: Immunoblotting of tIL-1RA protein expression in the different cell cultures. E-F: Representative confocal images of IL-1RA in different cell lines (D). IL-1RA localization in one immortal NOK cell line (E). Scale bar is 10 μm. G-H: Representative immunohistochemical pictures of IL-1RA expression in patient tissue samples. IL-1RA is downregulated in OD and OSCC samples, but occasionally, some OSCC retain some IL-1RA expression (G). Images were taken using a 10X objective (F). Mean expression scores of IL-1RA expression in the different groups (G). I-K: IL-1RN (I), icIL-1RN (J), sIL-1RN (K) mRNA expression in the different cell cultures according to their mortality state in culture. Expression is express as fold change relative to the reference gene. Data are shown as mean ± SEM (N = 3 independent experiments, n = 3 technical replicates). Two-tailed T-test was used to calculate the exact P value. Data information: *\*P < 0*.*05, **P < 0*.*005, P*** < 0*.*0007, P**** < 0*.*00001*.

In order to assess if OD and OSCC cells still retain the capability to upregulate IL1-RA under inflammatory stimuli, OD and OSCC cell lines with undetectable IL-1RA protein were treated with recombinant IL-1α or IL-1β (10 ng/ml). Neither IL-1α nor IL-1β affected tIL-1RA and icIL-1RN mRNA levels in OD and OSCC cell lines (Fig EV1C-F) but did induce IL-6 and IL-8 secretion, indicative of functional IL-1 signalling (Fig EV1G). tIL-1RA protein was not detected in any of the OD and OSCC cell lines before or after IL-1α or IL-1β stimulation (Fig EV1C-F). In order to assess whether another endogenous IL-1 inhibitor might be replacing this function in low-IL-1RA expressing cells, we analysed IL-1R2 expression, a decoy IL-1 receptor that acts as a molecular trap, inactivating IL-1 (Colotta et al., 1994). No difference in IL-1R2 mRNA levels between immortal NOK, immortal OD and OSCC cell lines were observed (Fig EV1H), and IL-1R2 transcript levels did not change after IL-1 stimulation in any of the tested cell lines (Fig EV1I). These data suggest that *IL1RN* downregulation in OD and OSCC cells is not reversible upon IL-1 stimulation and is not being compensated for by an increase in IL-1R2.

Immortal oral keratinocytes consistently expressed lower *IL1RN* transcript levels than their mortal counterparts (Fig EV1A, Fig 1I-K). Similarly, tIL-1RA protein levels decreased in immortal cells (Fig 1B). These data indicate a deregulation of the IL-1 inhibition system during oral carcinogenesis, which can occur at precancerous/dysplasia stages and which is more closely associated with the acquisition of replicative immortality than the stage of disease development (McGregor et al., 2002).

### IL-1R1 is upregulated in oral dysplastic and oral cancer cell lines compared to normal oral keratinocytes

To further characterize the regulation of IL-1 signalling during the oral carcinogenesis process, we analysed the expression of IL-1R1. IL-1R1 transcript levels were significantly up-regulated in immortal OD compared to iNOK cell lines (Fig 2A). Immunofluorescence showed endogenous expression of IL-1R1 in immortal OD and OSCC cells, but no expression in iNOKs (Fig 2B). IL-1R1 upregulation in OD and OSCC corresponded with higher endogenous levels of IL-1α protein (Fig 2C). In B16 (Fig 2D) and D20 (Fig 2E) cells, IL-1R1 was distributed diffusely across the cell membrane, but also inside or on the nucleus. To study the pattern of IL-1R1 expression under IL-1 stimulation, we treated B16 and D20 cells with recombinant IL-1α and recombinant IL-1β. IL-1α did not change IL-1R1 expression or localization, but treatment with IL-1β did increase IL-1R1 nuclear localization in both B16 and D20 cell lines (Fig 2 D, E). IL-1R1 nuclear expression increased significantly after IL-1β treatment in B16 cells (p = 0.0026) (Fig 2F).

**Figure 2:**
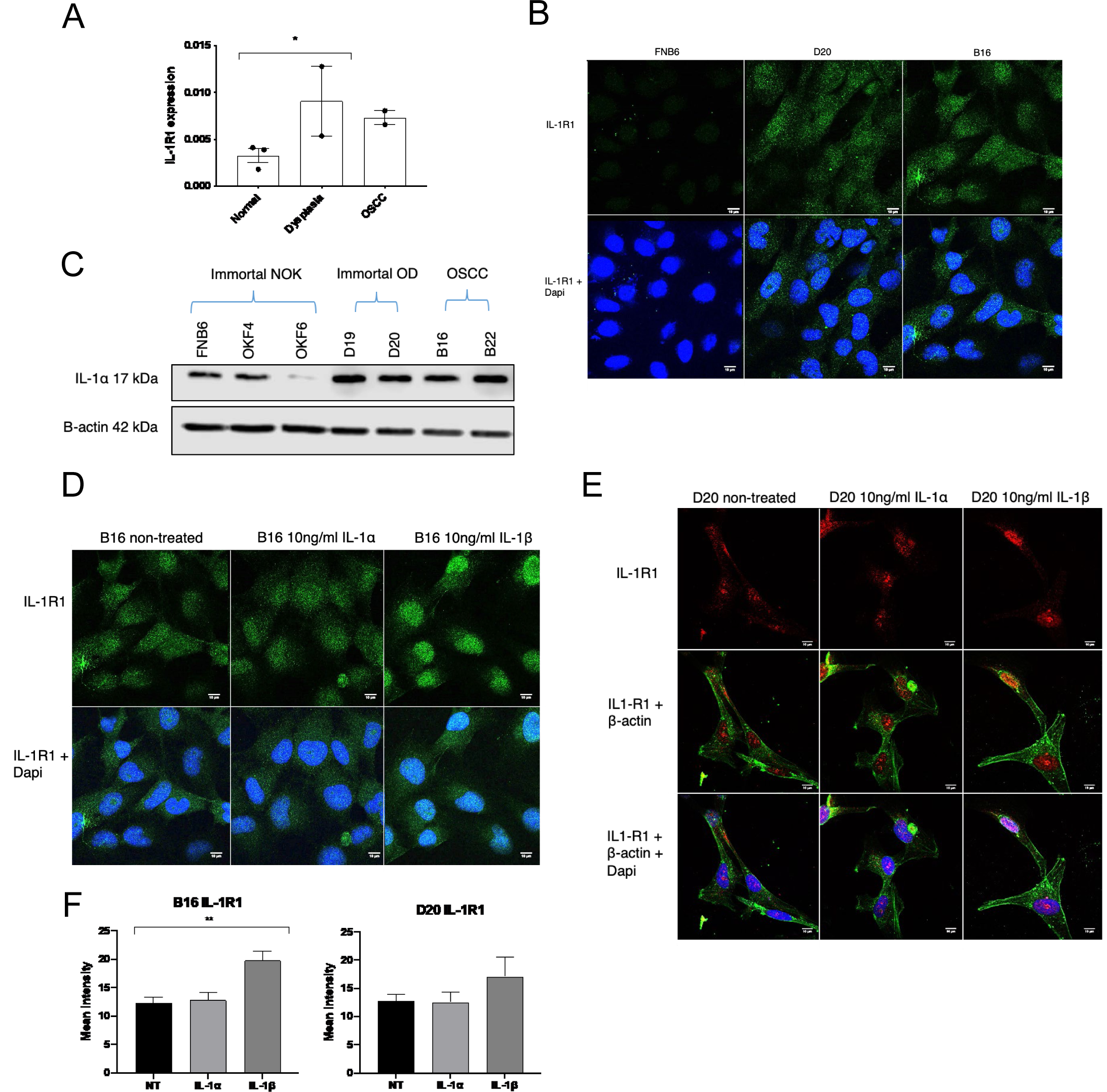
IL-1R1 is upregulated in oral dysplasia and oral cancer cell lines. A: IL-1R1 mRNA expression in immortal NOKs (FNB6, OKF4 and OKF6), immortal OD (D19 and D20) and OSCC (B16 and B22) cell clines. Expression is express as fold change relative to the reference gene. Data are shown as mean ± SEM (N = 3 independent experiments, n = 3 technical replicates). One-way ANOVA with multiple comparisons was used to calculate the exact P value. B: Representative confocal images of IL-1R1 expression in FNB6, D20 and B16 cell lines. Scale bars are 10 μm. C: Immunoblotting of IL-1α protein expression in the different immortal cell lines. D-E: Representative confocal images of IL-1R1 expression after stimulation with rIL-1α and rIL-1β in OSCC (D) and OD (E) cells. Scale bars are 10 μm. F: Quantification of IL-1R1 nuclear expression after stimulation with rIL-1α and rIL-1β in B16 and D20 cells. One-way ANOVA with multiple comparisons was used to calculate the exact P value (N = 2 independent experiments). Data information: *\*P < 0*.*05, **P < 0*.*005*.

### icIL-1RA1 regulates IL-6 and IL-8 secretion in mortal oral keratinocytes

Next, we knocked down (KD) icIL-1RA1 in primary normal and dysplastic oral keratinocytes (NOK805 and D6 respectively) using CRISPR/Cas9. As IL-1α is an important regulator of IL-6 and IL-8, we aimed to study if IL-1RA downregulation may favour oral cancer development by resulting in the overexpression of these cytokines. tIL-1RA protein levels in NOK805 KD (Fig 3A) and D6 KD (Fig 3B) cells were significantly lower than in their wild type (WT) counterpart. A low level of expression was retained, suggesting a heterogeneous population (Fig EV2A-D). Due to limitations in the replicative lifespan (8-10 passages), both cell types (KD and WT NOK805 and D6 cells) were maintained in culture with the Rho-kinase (ROCK) inhibitor (Y-27632), previously reported to delay the onset of senescence in primary keratinocytes (Chapman et al., 2014). When using the CRISPR-modified and control cells, Y-27632 was removed from the culture medium and senescence-associated β-galactosidase activity was assessed. This was done to confirm that the edited cells did not senesce immediately after removing Y-27632 (Fig EV3A, B) due to the drug-induced lifespan extension, and that the observed changes were attributed to icIL-1RA1 knock down and not due to the induction of senescence.

**Figure 3:**
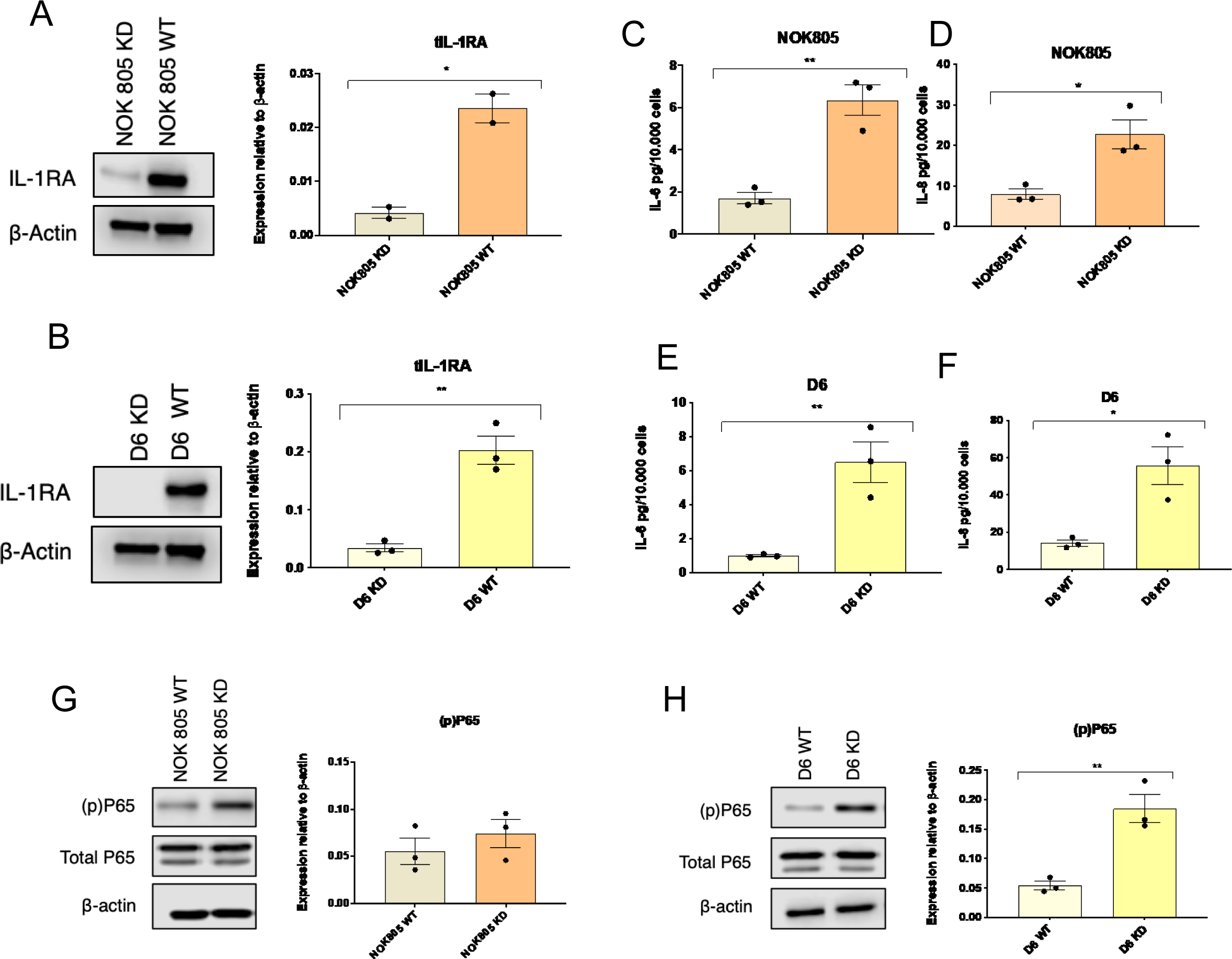
icIL-1RA1 regulates IL-6 and IL-8 secretion in normal and dysplastic mortal oral keratinocytes. A-B: icIL-1RA1 knock down in primary NOKs (A) and primary OD (B) after CRISPR/Cas9 gene edition. Quantification of tIL-1RA expression is expressed as mean ± SEM (N = 3 independent experiments). Two-tailed T-test was used to calculate the exact P value. C-F: IL-6 and IL-8 secretion by NOK805 cells (C, D) and D6 cells (E, F) after icIL-1RA1 gene edition. Data are shown as mean ± SEM (N = 3 independent experiments, n = 3 technical replicates). G-H: Immunoblotting of p65 phosphorylation after icIL-1RA1 gene edition in NOK805 (G) and D6 (H) cells. Quantification of (p)P65 expression is expressed as mean ± SEM (N = 3 independent experiments). Two-tailed T-test was used to calculate the exact P value. Data information: *\*P < 0*.*05, **P < 0*.*005*.

When comparing KD and WT cells, we found a significant increase in IL-6 and IL-8 secretion in both NOK805 (Fig 3C, D) and D6 (Fig 3E, F) KD cells compared to wild-type controls. To assess if these changes could be attributed to an increase in NF-κB pathway activation due to a lack of IL-1 inhibition, we analysed the activation NF-κB pathway by measuring the phosphorylation of p65. p65 phosphorylation increased in both NOK805 KD (Fig 3G) and D6 KD (Fig 3H) compared to WT cells, with no changes in total p65 levels. These data suggest that icIL-1RA1 modulates IL-6 and IL-8 levels by altering the activation of the NF-κB pathway in normal and dysplastic oral keratinocytes.

### icIL-1RA1 does not modify proliferation nor migration of immortal pre-malignant and malignant oral keratinocytes

To assess if re-expression of icIL-1RA1 in cells where IL-1RA expression has been lost could affect cell behaviour, we transfected two immortal cell lines with undetectable levels of IL-1RA protein (B16 and D20) with a vector encoding icIL-1RA1 (Fig 4A). Transfection efficiency was above 50% for both cell types (Fig 4B). Transfection of icIL-1RA1 did not affect proliferation (Fig 4C, D), nor alter cell migration (Fig E, F) in either cell line. In both B16 and D20 icIL-1RA1 transfected cells, a decrease in IL-6 and IL-8 secretion was observed, but only when cells were stimulated with IL-1β (Fig G-J).

**Figure 4:**
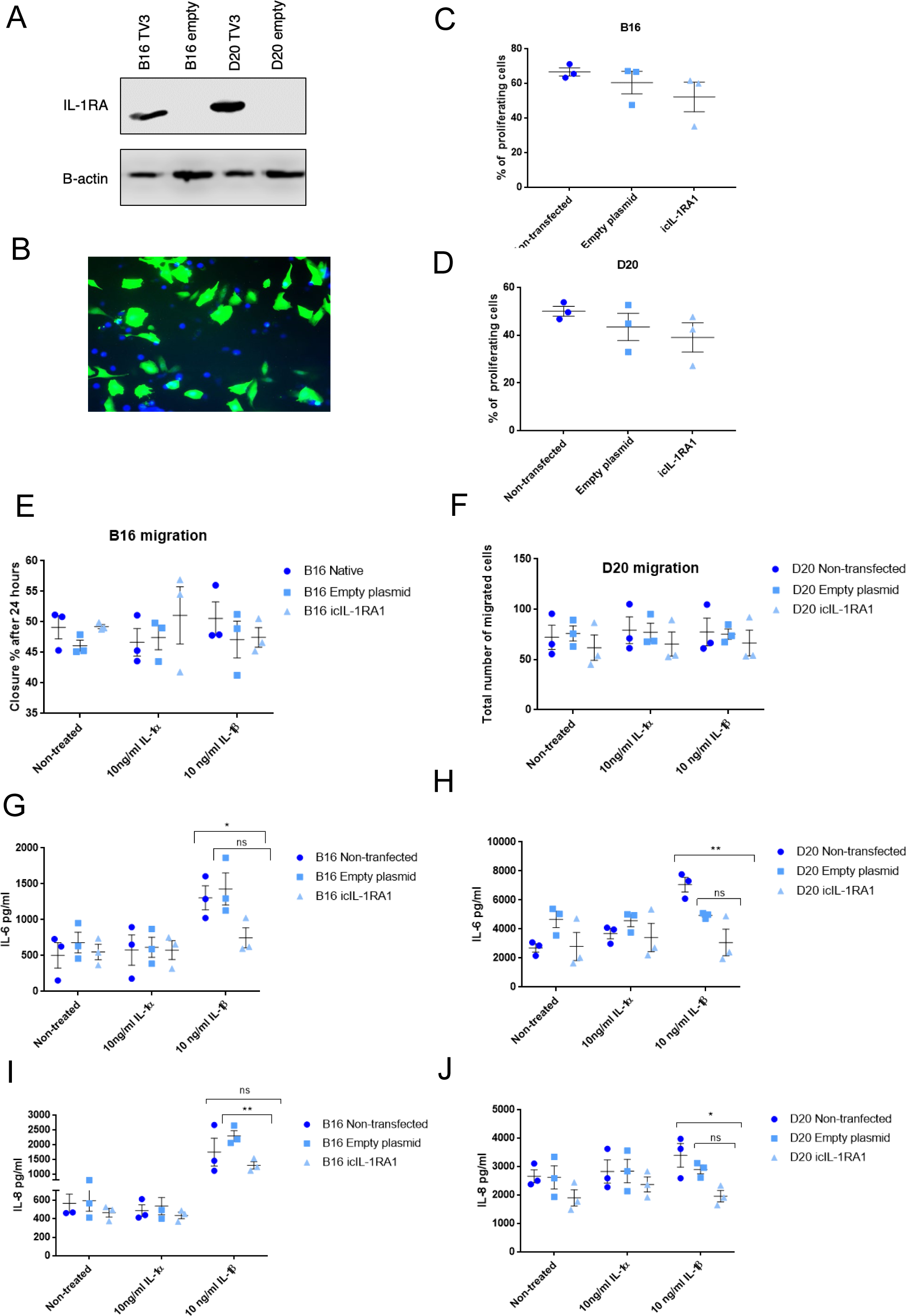
icIL-1RA1 does not modify proliferation nor migration of OD and OSCC cells. A-B: B16 and D20 cells were transfected with a pCMV6-Entry vector coding for icIL-1RA1 (A) with a transfection efficiency > 50% (B). C-D: Proliferation was assessed using the Click-iT™ Plus EdU Flow Cytometry Assay in icIL-RA1 transfected, mock transfected and non-transfected B16 (C) and D20 (D) cells. Data are shown as mean ± SEM (N = 3 independent experiments). One-way ANOVA was used to calculate the exact P value. E: The ORIS™ migration assay was used to assess B16 migration after transfection with a pCMV6-Entry vector coding for icIL-1RA1, an empty plasmid and non-transfected. Migration of each group was assessed with no treatment, after treatment with 10 ng/ml of rIL-1α and 10 ng/ml of rIL-1β. Data are expressed as mean area closure expressed as percentage ± SEM (N = 3 independent experiments, n = 3 technical repeats). Statistical analysis was done using two-way ANOVA. F: Migration of D20 cells after transfection was performed using the transwell system. Migration of each group (non-transfected, mock transfected and icIL-1RA1 transfected) was assessed with no treatment, after treatment with 10 ng/ml of rIL-1α and 10 ng/ml of rIL-1β. Data are expressed as mean number of migrated cells ± SEM (N = 3 independent experiments, n = 3 technical repeats). Statistical analysis was done using two-way ANOVA. G-J: IL-6 (G, H) and IL-8 (I, J) secretion after transfection with a pCMV6-Entry vector coding for icIL-1RA1, an empty plasmid and non-transfected. Secretion of both cytokines was assessed with no treatment, after treatment with 10 ng/ml of rIL-1α and 10 ng/ml of rIL-1β. Data are shown as mean ± SEM (N = 3 independent experiments, n = 3 technical repeats). Statistical analysis was done using two-way ANOVA. Data information: *\*P < 0*.*05, **P < 0*.*005, **** P < 0*.*00001*.

### IL-1RA expression decreases during replicative senescence

Replicative senescence is an important anti-tumour response, which paradoxically can also promote tumour formation if not tightly regulated. IL-1α has been identified as a key factor in the regulation of senescence and the SASP (Wiggins et al., 2019). Thus, we hypothesized that a decrease in tIL-1RA could switch the senescence response from preventing tumour formation, to promoting cancer development, by allowing an overexpression of inflammatory factors known to have pro-tumourigenic effects, such as IL-1, IL-6 and IL-8 (Acosta et al., 2013). In both normal (Fig 5A) and dysplastic (Fig 5B) oral keratinocytes, p16Ink4a, p21^Waf1/Cip1^ and SA-β galactosidase activity (Fig 5C) increased gradually with passage. Phosphorylation of H2AX (γH2AX) also increased significantly with time in culture of all cell types (Fig 5D-G).

**Figure 5:**
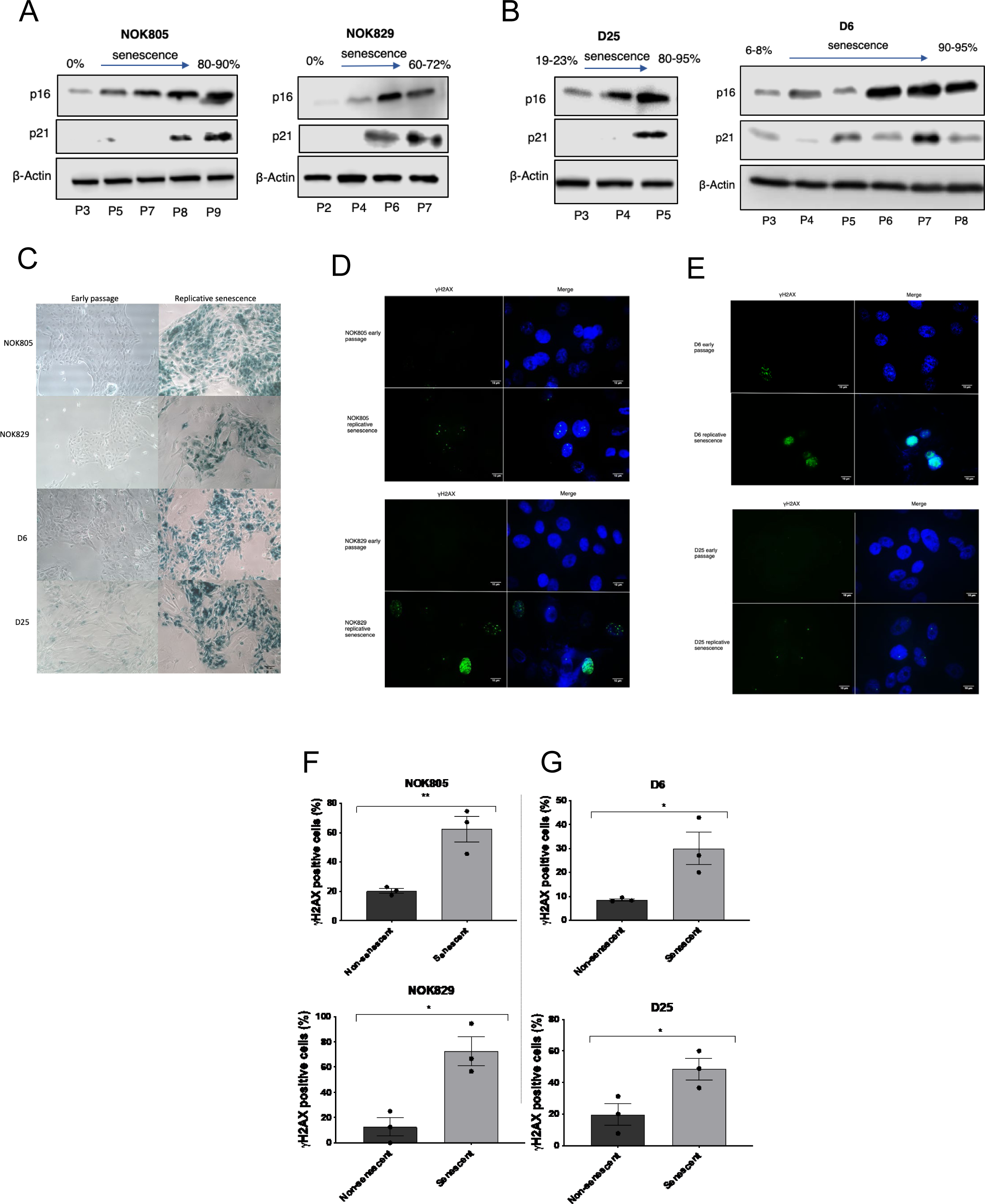
Senescence markers in normal and dysplastic oral keratinocytes. A-B: Immunoblotting showing p16^Ink4a^ and p21^Waf1/Cip1^ expression in normal (A) and dysplastic (B) oral keratinocytes during replicative senescence. C: Representative images of SA-β-GAL activity in non-senescent and senescent normal and dysplastic oral keratinocytes. Scale bar is 50 μm. D-G: Immunofluorescence and quantification of γH2AX expression in early passage and senescent normal (D, F) and dysplastic (E, G) oral keratinocytes. Data are shown as mean ± SEM (N = 3 independent experiments). Two-tailed T-test was used to calculate the exact P value. Scale bar is 10 μm. Data information: *\*P < 0*.*05, **P < 0*.*005*.

tIL-1RA protein expression decreased gradually with acquisition of replicative senescence in both normal (Fig 6A) and dysplastic cells (Fig 6C). icIL-1RN transcript levels also decreased significantly during aging in all of the cell cultures (Fig 6B, D). We then analysed the expression of commonly expressed SASP factors known to be able to promote tumour

formation: IL-1β, IL-1α, IL-6 and IL-8. Secretion of IL-1β increased in both senescent NOK (Fig 6E) and OD cell cultures (Fig 6F), and IL-1α protein expression increased in all senescent cell cultures apart from D6 cells (Fig 6G, H). Secretion of IL-6 and IL-8 (Fig 6I-L) and IL-6 and IL-8 mRNA transcript levels (Fig EV4A, B) increased gradually in all of the cell types during senescence. Interestingly, senescent ODs secreted more IL-6 and IL-8 than both senescent NOKs (Fig 6I-L) (apart from secreted IL-8 from NOK805 cells), suggesting an enhancement of the SASP in dysplastic cells.

**Figure 6:**
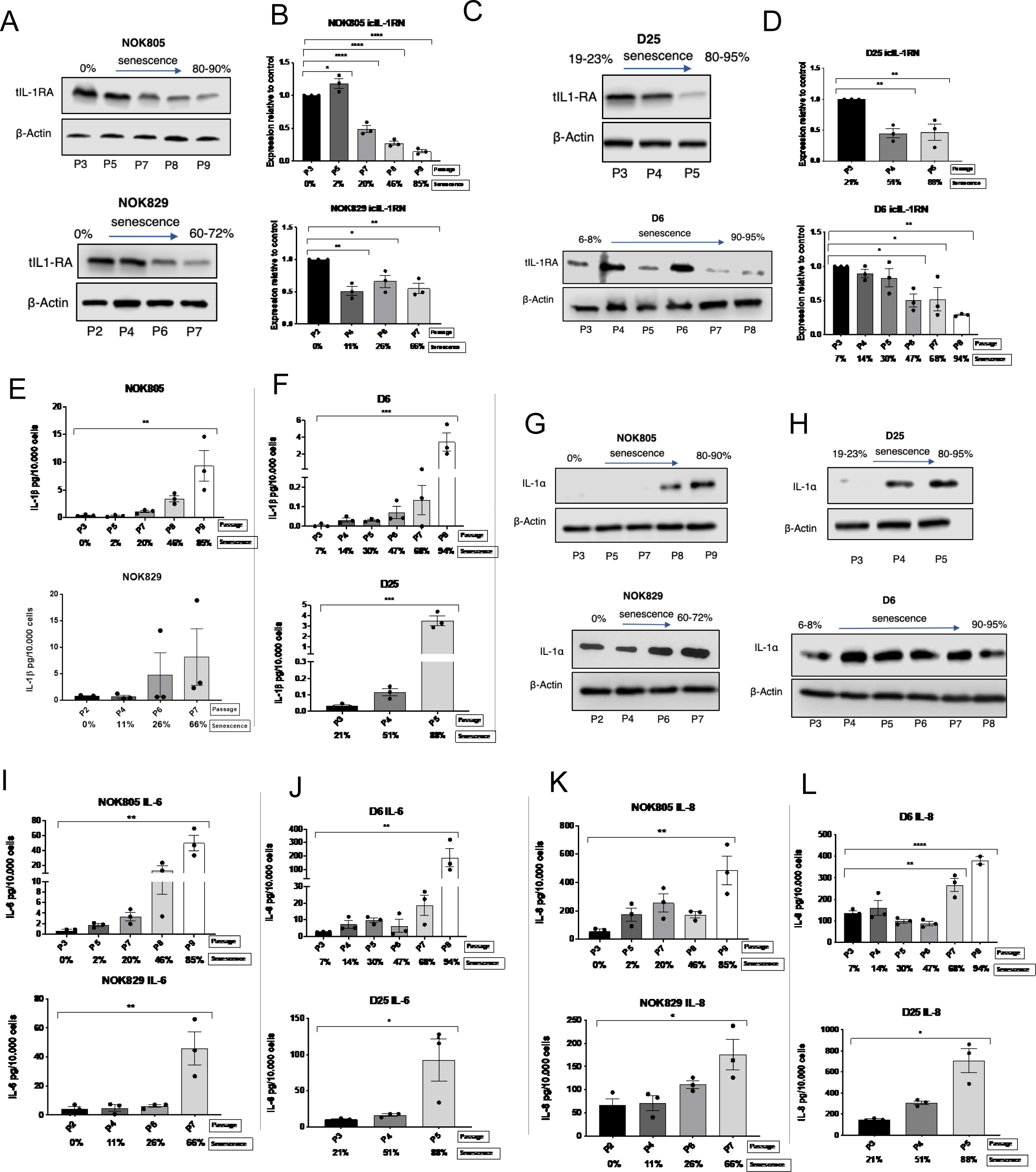
icIL-1RA expression decreases, while IL-6, IL-8, IL-1□ and IL-1 β levels increase during replicative senescence. A-D: Total IL-1RA protein expression and icIL-1RN transcript levels during replicative senescence of normal (A, B) and dysplastic (C, D) oral keratinocytes. E-F: IL-1β secretion during replicative senescence of normal (E) and dysplastic (F) oral keratinocytes. G-H: Immunoblotting showing IL-1α expression in normal (G) and dysplastic (H) oral keratinocytes during replicative senescence. I-L: IL-6 and IL-8 secretion during replicative senescence of normal (I, K respectively) and dysplastic (J, L respectively) oral keratinocytes. Data information: Data are shown as mean ± SEM (N = 3 independent experiments, n = 2 technical repeats). Statistical analysis was done using one-way ANOVA. *\*P < 0*.*05, **P < 0*.*005, ***P* ≤ *0*.*0005, **** P* ≤ *0*.*00001*

### The cGAS/STING pathway is activated during oral keratinocyte senescence

Senescent normal and dysplastic oral keratinocytes developed cytoplasmic chromatin fragments (CCF) (Fig 7A) probably as a consequence of a weakened nuclear membrane due to a decrease in Lamin B1 expression (Fig 7B). CCF have been reported to regulate senescence and the SASP by activating the cGAS/STING pathway (Dou et al., 2017, Gluck et al., 2017). Thus, we decided to assess activation of the cGAS/STING pathway during oral keratinocyte senescence. One of the downstream products of the cGAS/STING pathway is IFN-β. IFN-β mRNA transcript levels increased significantly in all cell types, except NOK805 cells, upon senescence (Fig 7C). To examine activation of the cGAS/STING pathway, we treated senescent dysplastic cells (D6) with the cGAS inhibitor, RU.521 (Vincent et al., 2017), and assessed changes in SASP components and NF-κB activation. Cell viability was assessed with trypan blue staining. RU.521 was not toxic at any of the used concentrations (Fig 7D). IL-6 secretion was significantly reduced by treating senescent cells with 10 µg/ml and 20 µg/ml of RU.521 (Fig 7E), but not with lower doses. Contrary to this, treatment with RU.521 had no effect on secreted IL-1β (Fig 7F) and IL-8 (Fig 7G). In accordance with the decrease in secreted IL-6, treatment with 10 µg/ml or 20 µg/ml of RU.521 also reduced phosphorylation of p65 (Fig 7H). A decrease in IL-1RA protein expression was also observed (Fig 7H), which may be a consequence of the decrease in p65 phosphorylation. A dose dependent increase of p16^Ink4a^ protein was also noted (Fig 7H).

**Figure 7:**
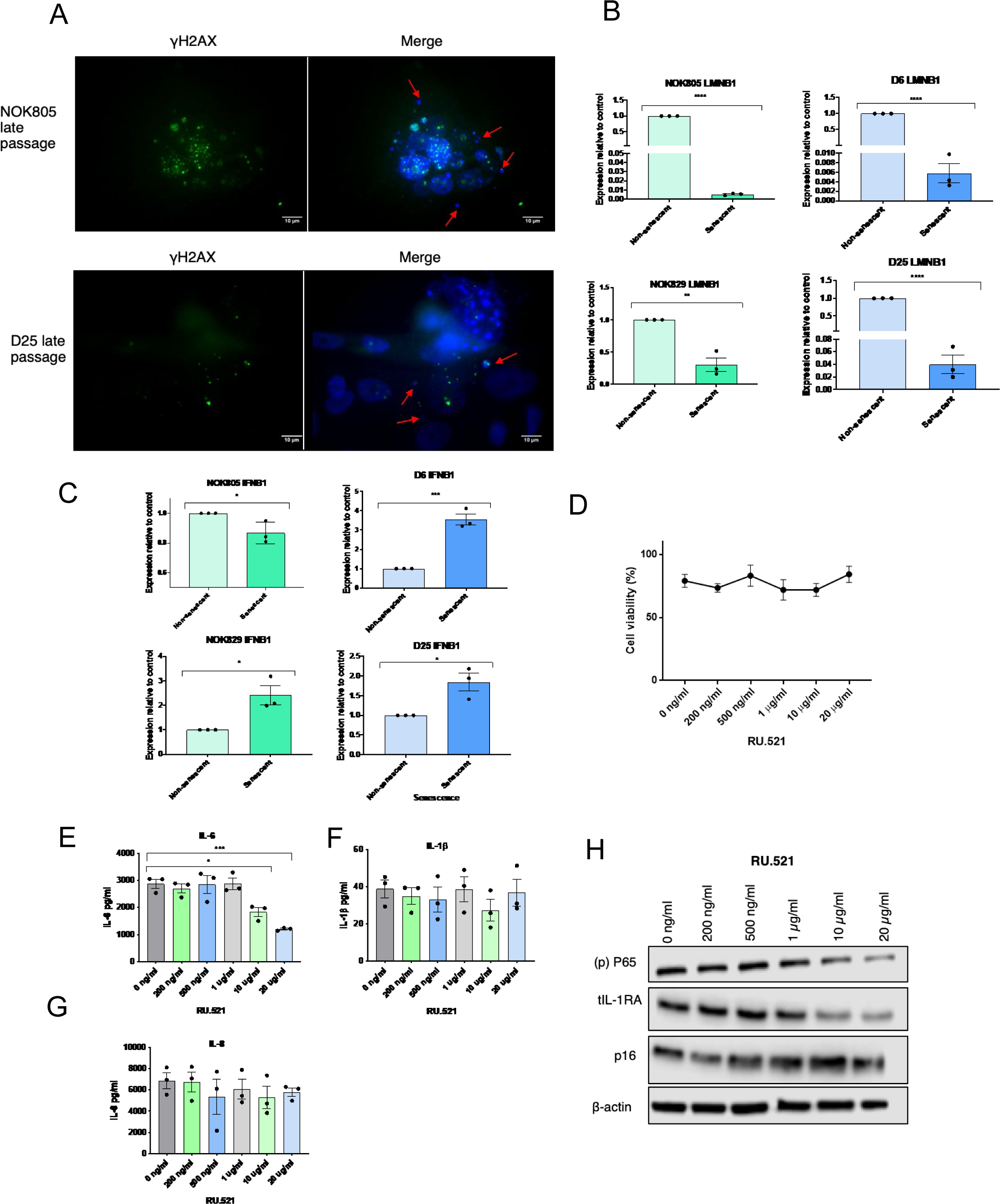
cGAS is activated during replicative senescence of oral keratinocytes. A: Immunofluorescence showing the presence of CCF in senescent NOK805 and D25 cells. Scale bar is 10 μm. B: Lamin B1 (LMNB1) transcript levels in senescent normal (NOK805 and NOK829) and dysplastic (D6 and D25) oral keratinocytes. Data are shown as mean ± SEM (N = 3 independent experiments, n = 3 technical repeats). Two-tailed T-test was used to calculate the exact P value. C: IFNB1 transcript levels in senescent normal and dysplastic oral keratinocytes. Data are shown as mean ± SEM (N = 3 independent experiments). Two-tailed T-test was used to calculate the exact P value. D: Cell viability of D6 cells after treatment with different doses of RU.521. Data are shown as mean ± SEM (N = 3 independent experiments). E-G: IL-6 (E), IL-1β (F) and IL-8 (G) secretion by D6 cells after treatment with RU.521 Data are shown as mean ± SEM (N = 3 independent experiments, n = 2 technical repeats). Statistical analysis was done using one-way ANOVA. H: Immunoblotting showing phosphorylation of P65, tIL-1RA and p16^Ink4a^ expression after treatment with RU.521. Data information: *\*P < 0*.*05, ***P < 0*.*0005*.

### icIL-1RA1 regulates the onset of senescence and the development of the SASP (Fig 8)

In order to study a specific role of icIL-1RA1 during senescence and the regulation of the SASP, we removed Y-27632 from WT and icIL-1RA1 KD D6 cells and expanded them until replicative senescence was achieved. Both KD and WT cells were at the same passage number when Y-27632 was removed from culture medium. D6 KD cells senesced prematurely compared to D6 WT cells (Figure 8A). On average, 87.5% of the D6 KD cells stained positive for SA-β-gal after one passage, compared with 27.9% of the D6 WT at the same PDs (Fig 8A, B). This corresponded with the cessation of cell growth observed in the KD cells, which could not be passaged further. Contrary to this, the D6 control cells were passaged one more time before achieving a SA-β-gal expression of 81% (Fig 8A, B). A statistically significant difference in population doublings before senescence was observed when comparing KD and WT cells (p < 0.0001): icIL-1RA1 KD cells doubled their population 0.7 times before senescing, while the WT cells doubled 2.4 times (Fig 8C).

**Figure 8:**
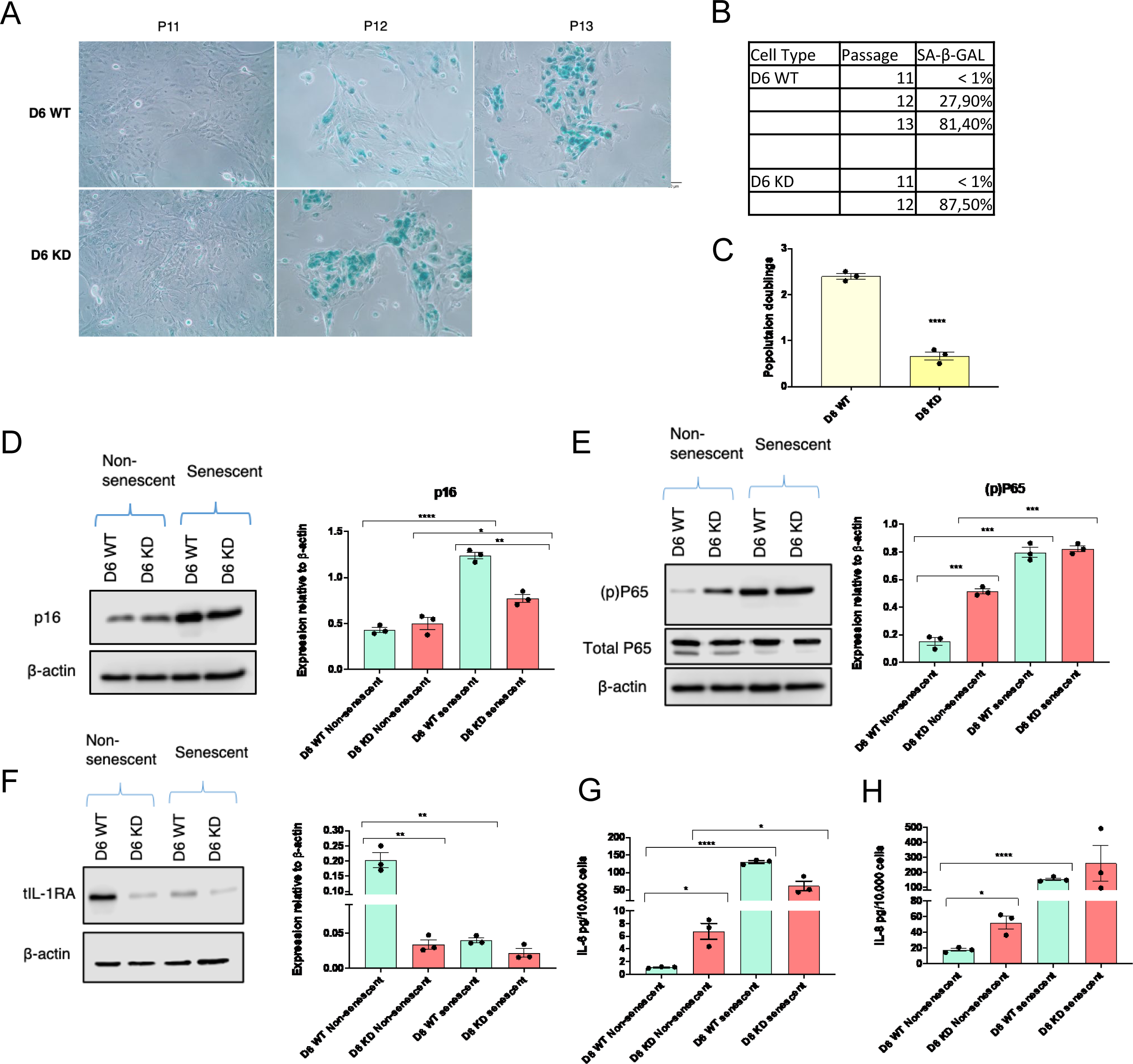
icIL1.RA1 regulates the onset of senescence and the SASP. A-B: SA-β-GAL activity after cell passaging in WT and icIL-1RA1 KD D6 cells. Scale bar is 50 μm. C: Comparison of population doublings between WT and icIL-1RA1 KD D6 cells. Data are shown as mean ± SEM (N = 3 independent experiments, n = 3 technical repeats). Two-tailed T-test was used to calculate the exact P value. D-F: Immunoblotting showing p16^Ink4a^ (D), phosphorylation of p65 (E) and tIL-1RA levels in non-senescent and senescent WT and icIL-1RA1 KD D6 cells. Data are shown as mean ± SEM (N = 3 independent experiments). Statistical analysis was done using one-way ANOVA. G-H: IL-6 and IL-8 secretion using same samples from D-F. Data are shown as mean ± SEM (N = 3 independent experiments, n = 2 technical repeats). Statistical analysis was done using one-way ANOVA. Data information: *\*P < 0*.*05, **P < 0*.*005, **** P < 0*.*00001*

Both D6 KD and WT cells showed an increase in p16^Ink4a^ protein expression following removal of Y-27632, with higher p16^Ink4a^ expression in WT than KD senescent cells (Fig 8D). As shown previously, knock out of icIL-1RA1 significantly increased phosphorylation of p65 (Figure 3G, H): this also further increased significantly in the senescent WT and KD cells (Fig 8E). Nevertheless, once the cells had senesced, both cell groups (KD and WT) exhibited a similar level of p65 phosphorylation. D6 KD cells expressed lower IL-1RA protein levels than D6 WT cells (this as a consequence of gene editing), but when cells had senesced, IL-1RA protein expression of WT cells decreased significantly, similar to the amounts observed in KD cells (Fig 8F). Once cells had senesced, both WT and KD groups experienced a significant increase in secreted IL-6 and IL-8. No difference in IL-6 and IL-8 secretion was observed between senescent WT and KD D6 cells (Fig 8G, H). Altogether, these findings suggest that in oral keratinocytes, icIL-1RA1 has a role in the onset of senescence and the development of the SASP, by regulating IL-6 and IL-8 levels through the NF-κB pathway.

## DISCUSSION

IL-1RA expression has been shown to decrease in oral squamous cell carcinomas (Shiiba et al., 2015). Evidence from other cancers suggest that exogenous IL-1RA might be effective for the treatment of IL-1 expressing tumours, such as melanoma, gastric and breast cancers, among others (Dinarello, 2010). A recent paper showed that exogenous IL-1RA can interrupt the oral carcinogenesis process *in vivo*, as submucosal injections of IL-1RA into the tongue of mice during 4NQO-induced oral carcinogenesis interrupted the malignant transformation process (Wu et al., 2016). Nevertheless, the timing, role and consequences of IL-1RA loss of expression during the malignant transformation of oral keratinocytes have not been explored in detail.

We found icIL-1RA was progressively downregulated in oral dysplastic (in both mortal and immortal, but more profoundly in immortal dysplasias) and oral carcinomatous cell lines, both at mRNA and protein level, but the mechanisms underlying the downregulation are not known. Post-transcriptional modifications such as methylation could be responsible for *IL1RN* downregulation in ODs and OSCCs, but only a small increase in methylation events of the IL-1RA promoter is evident in HNC (http://maplab.imppc.org/wanderer/), thus it is unlikely that this is the cause of loss of expression. *IL1RN* polymorphisms have been attributed to an increased risk of developing some cancers, like cervical gastric carcinomas (Wu et al., 2014, Zhang et al., 2012). Nevertheless, it is unlikely that this is the cause of loss of IL1-RN expression in OD and OSCC, as polymorphic genes are usually active and produce an abnormal form of the protein, which would not explain the decrease in *IL1RN* transcript levels and lack of protein expression in OD and OSCC cell lines. However, the mechanisms underlying *IL1RN* downregulation remain unclear. Other studies have also shown IL-1RA downregulation at mRNA and protein level in OSCC (von Biberstein et al., 1996, Shiiba et al., 2015, Hakelius et al., 2016). There has been little focus on when *IL-1RN* downregulation happens in the oral carcinogenesis process. As shown by our results, reduced expression of IL-1RA in dysplastic cells suggests that IL-1RA downregulation can occur in early (precancer) stages of the carcinogenesis process. This has also been reported by Shiiba et al. (2015), who found IL-1RA protein expression to be progressively downregulated through OD to OSCC.

There has been much discussion about the specific functions of icIL-1RA, and it has been hypothesized that its main function might be to counteract intracellular actions of IL-1α (Palmer et al., 2005). We found that IL-1RA was localized both in the cytoplasm and inside the nucleus of NOKs, which has also been reported in endothelial cells (Merhi-Soussi et al., 2005). Despite being immortal, the FNB6 cell line retained IL-1RA expression, possibly because it was immortalized at low passage number, where IL-1RA is high. Diffuse expression of IL-1RA in the cytoplasm of oral cells has been reported recently (Abe et al., 2017), but to our knowledge, we are the first group to report intra-nuclear IL-1RA expression in oral keratinocytes. This corresponds to our other finding that IL-1R1 also localizes both on the cell surface and inside or on the nucleus membrane of OD and OSCC cells. This is a novel finding, suggesting that intranuclear IL-1α (which is reported to act in a non-IL-1R1 dependent way) (Werman et al., 2004, Cheng et al., 2008) could also act in an IL-1R1 dependent manner, although that remains to be demonstrated. A recent report showed that full length IL-1α, which includes the nuclear localization sequence (NLS), is able to induce IL-1β, IL-6 and IL-8 secretion (Lau et al., 2019). IL-1R1 upregulation in immortal dysplastic and cancerous keratinocytes further highlights the de-regulation of the whole IL-1 system during oral cancer development, re-enforcing our hypothesis that IL-1RA downregulation is important for the acquisition of a malignant phenotype.

Contrary to previous reports from skin and bladder cancer cell lines (La et al., 2001, Merhi-Soussi et al., 2005), we did not find any effects of ectopic expression of icIL-1RA1 on migration nor cell proliferation. It is likely that the main advantage of icIL-1RA1 downregulation in oral epithelial cells is to allow an increase in IL-6 and IL-8 production, by enabling increased NF-κB activation, as observed when icIL-1RA1 was knocked down in mortal NOK and OD cell cultures. Both IL-6 and IL-8 are considered as “oncogenic cytokines”, as they are able to cause epithelial-to-mesenchymal transition (Coppe et al., 2008), stimulate angiogenesis and tumour growth (Sparmann and Bar-Sagi, 2004, Ancrile et al., 2007), disrupt cell-cell communication, impede macrophage function and promote epithelial and endothelial cell migration and invasion (Loaiza and Demaria, 2016). The activation of NF-κB in HNSCC has been reported to have an important role in the malignant phenotype of HNSCC, as inhibiting NF-κB inhibited growth, survival and expression of IL-1α, IL-6, IL-8 and GM-CSF in a mice head and neck cancer model (Duffey et al., 1999). In fibroblasts, the expression of IL-6 and IL-8 depends on NF-κB activation by the IL-1α/IL-1R1 axis, as IL-1α depletion has shown to significantly reduce IL-6 and IL-8 levels by reducing DNA binding activity of NF-κB in an IL-1R1 dependent manner (Orjalo et al., 2009).

As IL-1 is important for the development of the SASP(Lau et al., 2019), and there is evidence linking the SASP with malignant transformation of pre-cancerous cells (Malaquin et al., 2013, Coppe et al., 2008, Ortiz-Montero et al., 2017, Lau et al., 2019), we decided to study the role of icIL-1RA during senescence of normal and dysplastic oral keratinocytes. icIL-1RA1 depletion in D6 cells increased the secreted levels of IL-6 and IL-8 in pre-senescent cells and triggered premature senescence. In agreement with this, Uekawa et al., (2004) showed that depletion of IL-1RA in mouse embryonic fibroblasts (MEFs) triggered premature senescence, probably via activation of the p38 MAPK by IL-1β. Also, IL-1β has also been reported to induce senescence in NOKs (Jang da et al., 2015) and IL-1α has been shown to regulate the induction of senescence in a paracrine manner (Acosta et al., 2013). This suggests that icIL-1RA is an indirect regulator of the onset of senescence, by regulating IL-1 activity, which is important for the induction of senescence (Fig 9A). Nevertheless, according to a recent paper from Lau et al. (2019), both IL-1α and IL-1β are crucial for the development of the SASP in an IL-1R1 dependent manner, but are not essential for the triggering of senescence, as cells in which IL-1α and IL-1β have been inactivated still senesce (Lau et al., 2019).

**Figure 9:**
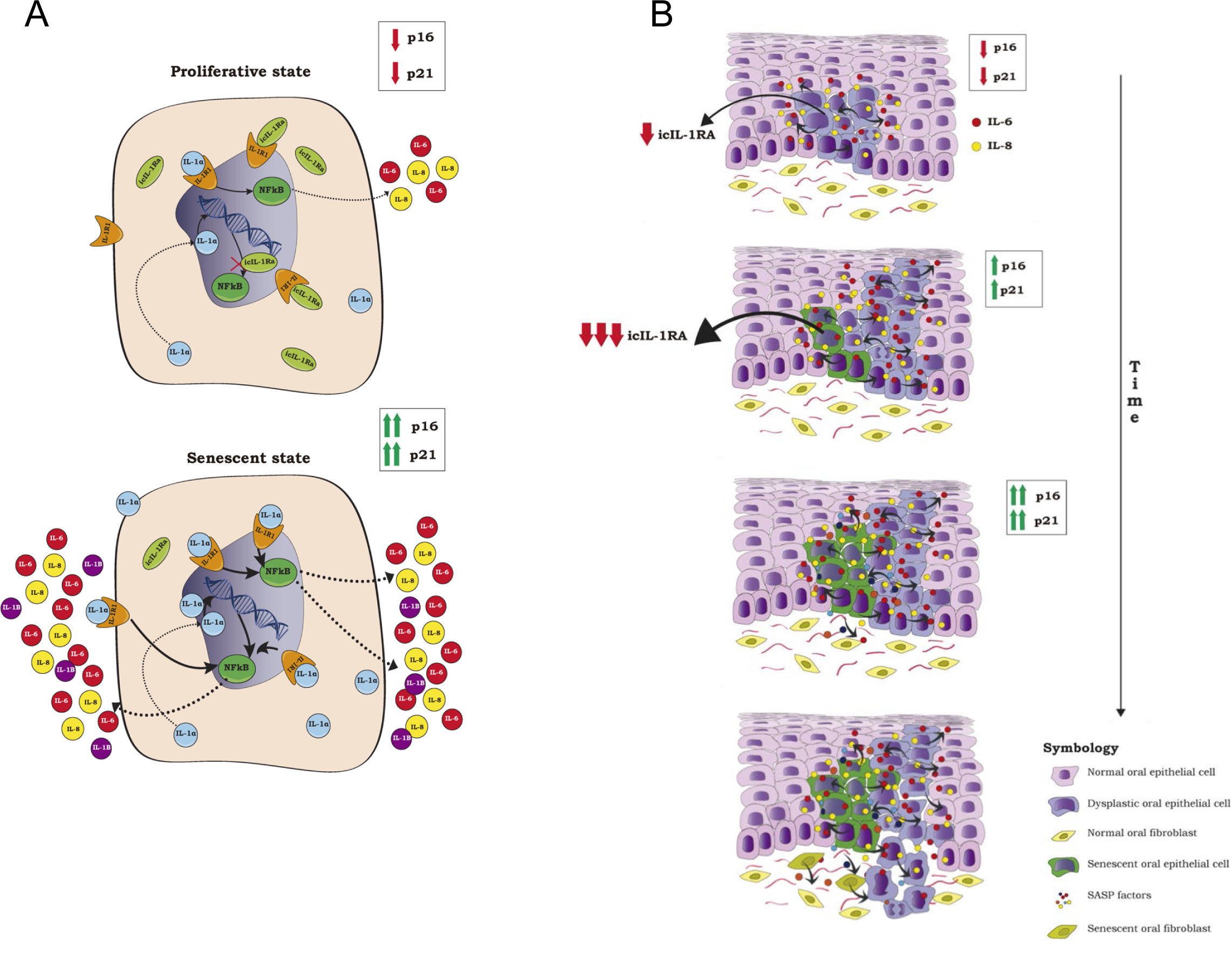
A proposed model of how icIL-1RA1 might regulate senescence and the SASP and how this may be related to oral cancer development. A: If icIL-1Ra1 expression decreases, IL-1α can bind to more IL-1R1 receptor and intranuclear IL-1α can interact with the nucleus with no or less regulation, which would increase the activation of the NF-κB pathway. This will lead to the production of more IL-1α, IL-1β (IL-1β is not constitutively produced by oral keratinocytes) and to high levels of IL-6 and IL-8, which will induce and re-enforce the senescent state. B: Downregulation of icIL-1RA1 in dysplastic oral keratinocytes facilitates an increase secretion of IL-6 and IL-8, which can promote EMT in dysplastic cells if left over time. In addition, many dysplastic keratinocytes will senesce due to a DDR, but some of them will escape senescence, probably due to the acquisition of mutations enabling immortality. So, if the dysplastic cells are not removed or the inflammatory microenvironment is not regulated, the SASP can act as a promoter on the “initiated” immortal cells, inducing malignant transformation.

We made the observation that cytoplasmic chromatin fragments (CCFs) were present in both normal and dysplastic senescent oral keratinocytes. This was correlated with a significant decrease in Lamin B1 expression, which suggested an increase in nuclear membrane permeability. CCF can initiate senescence by triggering the innate immunity cytosolic DNA-sensing cGAS-STING pathway (Dou et al., 2017). We inhibited cGAS in senescent ODs (D6 cell culture) with a cGAS inhibitor, RU.521 (Vincent et al., 2017). RU.521 significantly decreased secreted IL-6 levels when used at a concentration of 10 µg/ml and 20 µg/ml, with no changes observed in secretion of IL-8 and IL-1β. Similar to these results, cGAS has been reported to regulate the SASP by decreasing IL-6 levels *in vitro* and *in vivo* (Gluck et al., 2017). Interestingly, we observed a dose dependent decrease in phosphorylated p65 and IL-1RA proteins levels, indicating a decrease in NF-κB activity. p16 protein expression rose in a dose dependent manner. This was unexpected, as removal of cGAS has been shown to prevent the onset of senescence and decrease p16^Ink4a^ expression (Gluck et al., 2017). One possible explanation is that in this experiment cGAS was inhibited in already senescent cells, where p16^Ink4a^ regulation might be different. There was no evidence that the effects of RU.521 arose as a consequence of cytotoxicity, as cell viability did not change between untreated and treated groups. These results indicate that the cGAS/STING pathway is activated during replicative senescence of oral keratinocytes, which has not been reported before. It also suggests that there is feedback regulation between the NF-κB pathway and IL-1RA, as knock out of IL-1RA resulted in an increase in p65 phosphorylation, whereas a decrease in p65 phosphorylation (by cGAS inhibition) resulted in a decrease in IL-1RA.

Altogether our data suggests that icIL-1RA1 downregulation can contribute to oral cancer development in two ways; by facilitating a pro-inflammatory state in non-senescent cells and by allowing a premature and deregulated SASP, which are likely to be acting in combination (Fig 9A). icIL-1RA1 expression is lost early in the oral carcinogenesis process, often at the dysplasia stage, particularly seen when dysplastic cells achieve replicative immortality. This facilitates an increase in the activation of the NF-κB pathway which supports the development of a pro-inflammatory state, with high levels of IL-6 and IL-8. If dysplastic cells (in which icIL-1RA1 expression is lost or decreased) are not removed overtime, an inflammatory microenvironment will be generated. In vivo, it is very common to observe inflammatory infiltrate below the areas of epithelial dysplasia (Napier and Speight, 2008). Nevertheless, icIL-1RA1 downregulation results in an increase in IL-6 and IL-8, and both of which can promote EMT (Coppe et al., 2008) and IL-6 can act as a mitogen in a paracrine fashion (Kuilman et al., 2008). In addition, inflammation has been shown to contribute to OSCC invasion (Goertzen et al., 2018). Thus, if the dysplastic cells with low levels of icIL-1RA1 are not eliminated or the inflammatory response is not regulated, the imbalance in the inflammatory response could promote the development of an OSCC. In addition, oral dysplasias are a mixture of mortal and immortal cells (McGregor et al., 2002). Many mortal dysplastic cells will senesce as part of the DDR, and normal keratinocytes will senesce as a consequence of replicative ageing. This will lead to the development of the SASP (which is overexpressed in OD compared to NOKs). This should be beneficial as it could prevent dysplastic cells from progressing into cancer. But not all dysplastic cells will senesce. Some will escape senescence, e.g. because of p16^Ink4a^ mutation or methylation, mutations or inactivation of p53 and reactivation of telomerase and will become immortal (McGregor et al., 2002, McGregor et al., 1997). Thus, the “initiated” immortal cells are primed for malignant transformation, on which the SASP can act as a “promoter” (Fig 9A) (Coppe et al., 2008, Ortiz-Montero et al., 2017). Supporting this idea is the fact that *in vivo*, all leukoplakias have some degree of p16^Ink4a^ expression which is not related to HPV infection (Tomo et al., 2020) and the presence of γH2AX foci in oral dysplasia is related with the risk of malignant transformation, and although γH2AX is a marker of genomic damage/instability, it does also mark senescent cells (Leung et al., 2017).

Our findings suggest that targeting the IL-1 pathway by restabilising IL-1RA levels in oral dysplasias could be a useful approach to reduce malignant transformation of OPMDs.

## MATERIALS AND METHODS

### Cell culture

Isolation of primary normal oral keratinocytes (NOK) was done under The University of Sheffield ethics (approval reference 3463). The following cell lines/cell cultures used in this study have been described previously: FNB6, D6, D25, B16, B22, D19, D20 (McGregor et al., 1997, McGregor et al., 2002), OKF4 (Rheinwald et al., 2002), OKF6 (Natarajan et al., 2006).

FNB6, OKF4, OKF6, B16, B22, D19, D20 cell lines were cultured in Dulbecco’s Modified Eagle Medium low glucose (DMEM) (Sigma-Aldrich, Gillingham, UK) containing 10% vol/vol foetal bovine serum and 21% vol/vol F12 nutrient mix (Sigma-Aldrich, Gillingham, UK), supplemented with 2 mM L-glutamine (Sigma-Aldrich, Gillingham, UK), 6.25 μg/ml adenine, 100 μg/ml penicillin and 100 U/ml streptomycin (Sigma-Aldrich, Gillingham, UK), 8.47 ng/ml cholera toxin, 10 ng/ml epidermal growth factor (EGF) (Sigma-Aldrich, Gillingham, UK) and 5 ug/ml human insulin (Sigma-Aldrich, Gillingham, UK). D6, D25, NOK805 and NOK829 cell cultures were grown using lethally irradiated (i) 3T3 feeders as described previously (Rheinwald and Green, 1975) in Dulbecco’s Modified Eagle Medium low glucose (DMEM) (Sigma-Aldrich, Gillingham, UK) containing 10% vol/vol foetal bovine serum (Hyclone FetalClone II Serum, Fischer Scientific, New Hampshire, US) and 21% vol/vol F12 nutrient mix (Sigma-Aldrich, Gillingham, UK), supplemented with 2 mM L-glutamine (Sigma-Aldrich, Gillingham, UK), 0.25 μg/ml adenine, 100 μg/ml penicillin and 100 U/ml streptomycin (Sigma-Aldrich, Gillingham, UK),10 ng/ml epidermal growth factor (EGF) (Sigma-Aldrich, Gillingham, UK), 1 mg/ml human insulin (Sigma-Aldrich, Gillingham, UK), 0.4 μg/ml hydrocortisone, 1.36 ng/ml and 5 μg/ml 3, 3, 5-Tri-iodothyronine/ Apo-Transferrin and 0.7 mM Na Pyruvate (Sigma-Aldrich, Gillingham, UK). All cell cultures were incubated at 37°C with 5% CO2 and checked daily. Medium was replaced every 2-3 days and cells were passaged when approximately 80-90% of confluency was achieved. All cell cultures were routinely tested for mycoplasma infection.

### Confirmation of identity-STR profiling

DNA was extracted from each cell type with the Wizard Genomic DNA Purification Kit (Promega, Southampton, UK) and send for STR profiling (Core Genomic Facility, Medical School, The University of Sheffield). No match was found for any of the cells tested, which was expected as none of the cell lines used had been previously registered, but the results were comparable to previous analysis. Lack of match also shows that contamination with other cell lines was unlikely.

### Transient transfection

Approximately 2×10^5^ cells per well were seeded in 6-well plates in growth media free of antibiotics and were left at 37 °C and 5% CO2 for 24 hours. The next day, cells were approximately 70-80% confluent and were transfected using FuGENE HD transfection reagent (Promega, Southampton, UK) following the manufacturer’s recommendations. Cells were whether transfected with a pCMV6-Entry vector coding for icIL-1RA1 or a pCDNA3.1 empty vector (Origene, Rockville, USA) and incubated at 37 °C and 5% CO2 for 48 hours. After 48 hours, cells were ready for experimentation and/or analysis. After every transfection procedure, immunoblotting was done using an anti-IL-1RA antibody, in order to confirm the presence of the transfected protein.

### icIL-1RA1 knock down

Intracellular IL-1RA1 knock down was performed using the CRISPR/Cas9 system based on the protocol from Ran et al. (2013). Two commercially available sgRNAs (1 and 2) targeting a different segment of 20 nucleotides of exon 3 of the icIL-1RA1 gene were bought already incorporated into a pSpCas9 BB-2A-GFP vector (PX458) from GeneScript (Leiden, Netherlands). The sgRNA sequences are as follows: sgRNA1 F: 5’ TCCTCAGATAGAAGGTCTTC 3‘, R: 5’ GAAGACCTTCTATCTGAGGA 3’, sgRNA2 F: 5’ CTAGTTGCTGGATACTTGCA 3‘, 5’ TGCAAGTATCCAGCAACTAG 3’. NOK805 and D6 cells were seeded into 6-well plates at a density of 2,5 × 10^5^ cells per well in antibiotic free media with 10 μg/ml Y-27632 (Abcam, Cambridge, UK) and left to adhere at 37 °C and 5% CO2. Y-27632 (Abcam, Cambridge, UK) was used in order to facilitate cell growth and to avoid the induction of senescence. After 24 hours, cells were transfected using JetPrime (Polyplus, Illkirch, France) following the manufacturer’s recommendations. Twenty-four hours after transfection, cells were re-suspended in growth media and sorted using a FACSAria IIu cell sorter (BD biosciences, San Jose, USA). The machine was previously calibrated with non-transfected cells of each cell type and baseline for fluorescence was set. Cells were divided into GFP positive (+) and GFP negative (-) cells and same number of cells were collected per group. The gate was adjusted strictly in order to diminish the chance of getting any false positives into the GFP+ group. GFP-cells were used as controls for all downstream assays. Single colony expansion could not be done as both cell types used (NOK805 and D6 cell cultures) did not grow in isolation. GFP-and GFP+ cells were grown under the same conditions and analysed or used at the same passage number.

Genomic GNA (gDNA) was extracted from each of the GFP+ and GFP-cell types using the Wizard Genomic DNA Purification Kit (Promega, Southampton, UK) and quantified using a Nanodrop 1000 Spectrophotometer (Thermo Fischer Scientific, Cambridge, UK). Primers targeting a sequence surrounding and including both sgRNA target sites were designed (F: 5’ GCTGGGCACATGGTGGCTGT 3’, R: 5’ ATGCCCACATACATTGGCTT 3’) and PCR reactions for each of the gDNAs were performed as follows: 5 minutes at 95°C and 30 cycles of 95 °C for 60 seconds, 55 °C for 60 seconds, 72 °C for 30 seconds followed by 2 minutes at 72 °C. PCR products were run in a 2% agarose gel for 90 minutes at 90 V and visualized using the InGenius3 gel documentation system (Syngene, Bangalore, India). PCR bands were extracted from the agarose gel and purified using and Isolate II PCR and gel kit (Bioline, London, UK) and sent for Sanger sequencing (GATC Biotech, Eurofins Genomics). Sequences obtained from WT cells were aligned with the expected sequence in order to corroborate proper primer amplification. This was done using the multiple sequence alignment tool of the European BioInformatics Institute (EMBL-EBI) (https://www.ebi.ac.uk/Tools/msa/clustalo/). In order to confirm a successful editing event, the sequence obtained from WT and CRISPR edited cells were aligned together.

### Senescence assays

Normal oral keratinocytes (NOK805 and NOK829 cell cultures) and mortal oral dysplasias (D6 and D25 cell cultures) were grown in combination with i3T3 in T25 cm2 tissue culture flasks until replicative senescence was achieved. Cells were passaged when they achieved a confluency of 70-85%. Before trypsinization, conditioned media was stored at -20°C and i3T3 feeders were removed by incubating them with 0.02% EDTA solution for 3-5 minutes at 37 °C and 5% CO2. After every passage senescence associated β galactosidase activity was quantified.

### Senescence associated β galactosidase staining

Twenty thousand cells per well were seeded in a 12 well-plate and left to adhere overnight. Senescence associated β-Galactosidase activity was assessed using a senescence detection kit (ab65351, Abcam, Cambridge, UK) following the manufacturer’s recommendations.

### Proliferation assay

Proliferation was assessed with the Click-iT(tm) Plus EdU Flow Cytometry Assay (Invitrogen, Cambridge, UK) following the manufacturer’s recommendations.

### Migration assays

To assess migration properly, cell proliferation was inhibited with Mitomycin C. Mitomycin C concentration was optimised using plots of cell replication (CellTrace(tm), Invitrogen, Cambridge, UK) through flow cytometry. Concentrations of 0.5 ug/ ml and 2 ug/ ml were used to inhibit cell growth from B16 and D20 cell lines respectively.

For the B16 cell line, migration was assessed using the ORIS(tm) cell migration assay (Platypus Technologies, Madison, USA) following the manufacturers recommendations. For the D20 cell lines, the transwell migration assay was used, as those cells were not suitable for the ORIS(tm) cell migration assay. Before seeding the cells, the external surface of the membrane (PET membrane, 8.0 µm, 24-well inserts, Falcon, Loughborough, UK) was coated with fibronectin. This was done by adding 100 µl of 10 µg/ml fibronectin (Sigma-Aldrich, Gillingham, UK) and incubating each insert for 1 hr at 37 °C. After this time, 100 µl and 300 µl of 1% BSA serum-free media were added inside the insert and on the bottom well, correspondingly. Each plate was incubated for 1 hr at 37 °C. This was done in order block non-specific binding. Once the inserts were properly coated with fibronectin, cells were trypsinized, centrifuged at 1,000 rpm for 5 minutes and re-suspended in 1% BSA serum-free media. Cell suspension was further diluted in order to achieve the desired cell concentration. In-between, 500 µl of 10% FBS containing media and mitomycin C at a concentration of 2 µg/ml were added. Once this was done, 100 µl of cell suspension containing approximately 25,000 cells were added into each insert and incubated for 3.5 hours at 37 °C and 5% CO2 After this, the media from each bottom well was removed (in order to remove the mitomycin C) and 500 µl of 10% FBS containing media with rIL-1α (10 ng/ml), rIL-1β (10 ng/ml) or nothing extra was added to each bottom well and incubated for 24 hours at 37 °C and 5% CO2. The next day, media was removed, each insert was washed once in PBS and fixed for at least 20 minutes in 10% formalin. The inserts were then washed once in distilled water and stained for 30 minutes in a solution of 0.5% (w/v) of crystal violet in 10% ethanol. Following this, the inserts were washed 2-3 times in distilled water before proceeding to the mounting stage. The cells attached to the inner part of the membrane (top), were swabbed away using a cotton bud, in order to keep just the cells that migrated from the inner side (top) into the outer side (bottom) of the membrane. The membranes were then removed from the insert by cutting them carefully with a scalpel blade and were then mounted on glass slides using DPX mounting media (Merck). Three pictures of each experimental membrane were taken using an Olympus BX51 optical microscope at 20X magnification, and cells were counted using Fiji ImageJ.

### cGAS inhibition assay

Mortal dysplastic keratinocytes (D6) were grown until replicative senescence was achieved (evaluated by SA-β-Gal activity). A total of 160 x10^4^ senescent cells were seeded in 12-well-plates and left overnight to adhere. D6 senescent cells were then treated for a period of 24 hours with different concentrations of the cGAS inhibitor RU.521 (0 ng/ml, 200 ng/ml, 500 ng/ml, 1µg/ml, 10 µg/ml and 20 µg/ml) (InvivoGen, California, USA) After 24 hours incubation at 37 °C and 5% CO2, conditioned media was collected and cells were harvested for analysis. Each individual repeat was counted, and the proportion of live/dead cells was calculated for each experimental group. This was done by staining the cells with Trypan blue (Thermo Fischer Scientific, Cambridge, UK) and counting them using a haemocytometer.

### Immunofluorescence

Between 20,000 and 40,000 cells were seeded on 8-well tissue culture chambers (Starstedt, Nümbrecht, Germany) and incubated for 24 hours at 37 °C and 5% CO2. Cells were then washed three times with PBS and fixed with 4% para-formaldehyde (Santa Cruz Biotechnologies, Texas, US) for 10 minutes. After fixation, cells were washed three times with PBS and blocked for 1 hour with 0.2% triton X-100 and 3% BSA (Sigma-Aldrich, Gillingham, UK) in PBS. After blocking, the cells were washed three times with PBS and incubated with primary antibody at working concentration in 3% BSA for 1-2 hours. Primary antibodies used were: goat anti-IL-1RA IgG (1: 500, AF-280-NA, R&D Systems, Minneapolis, USA), mouse anti-γH2AX IgG (1:200, JBW301, Merk Millipore, Massachusetts, USA). Negative controls were incubated without the primary antibody under the same conditions. Samples were then washed three times with PBS and incubated with conjugated secondary antibody in 3% BSA for 1 hour under darkness. Secondary antibodies used were: Alexa Fluor 594 donkey anti-goat (1: 1000, A11058, Thermo Fischer Scientific, Cambridge, UK), Alexa Fluor 488 goat anti-mouse (1: 1000, A11001, Thermo Fischer Scientific, Cambridge, UK). After secondary antibody incubation, the samples were washed 3 times with PBS followed by 3 washes with distilled H2O and were mounted using Prolong (tm) Gold antifade mountant with Dapi (Invitrogen, Cambridge, UK). The slides were left to set for 24 hours in the dark at 4°C before visualization. Samples were visualized using a Zeiss 880 AiryScan confocal microscope (Carl Zeiss, Oberkochen, Germany) or an Axioplan 2 Fluorescent microscope (Carl Zeiss, Oberkochen, Germany). Images were then processed with Fiji-ImageJ. Negatives controls of each individual cell line were processed together with each sample using the same settings.

For double labelling, cells were blocked in a solution of 10% Goat serum (the species in which the first secondary antibody was raised) with 0.2% triton X-100 in PBS. The samples were then incubated with the first primary antibody (an anti-IL-1R1 antibody, 1: 100, MAB269, R&D Systems, Minneapolis, USA). Negative controls were incubated in the same solution without the primary antibody. The secondary antibody used was a goat anti mouse Alexa fluor 594 IgG (ThermoFisher Scientific, Cambridge, UK), at a concentration of 1: 1000. After incubation with the first secondary antibody for 1 hour, the samples were washed 3 times with PBS and then blocked with a solution of 10% rat serum, 0.2% triton X-100 in PBS for 60 minutes. After this, the cells were washed three times with PBS and were incubated with the second primary antibody, an anti β-actin antibody (A1978, Sigma-Aldrich, Gillingham, UK) at a concentration of 1: 200 in a solution of 3% BSA serum, 0.2% triton X-100 in PBS for 1 hour. Negative controls were incubated in the same solution without the primary antibody. After this, the samples were washed three times with PBS and incubated in the second secondary antibody, a goat anti mouse Alexa fluor 488 IgG (H+L) (Thermofisher Scientific, Cambridge, UK) (1: 1000) in a solution of 3% BSA serum, 0.2% triton X-100 in PBS for 1 hour. Next steps were done as stated above.

### Immunohistochemistry

IHC for IL-1RA was performed on 4 μm formalin-fixed paraffin-embedded sections. Human skin sections were used as positive controls. Antigen retrieval was performed with heat-induced epitope retrieval in sodium citrate buffer (10 mM sodium citrate, 0.05% Tween 20, pH 6.0). Primary antibody used was goat IgG anti-IL-1RA (1: 500, AF-280-NA, R&D Systems) and samples were incubated overnight at 4°C. After secondary antibody incubation, staining was visualized with the Vectastain ABC Kit (Vector Laboratories, Burlingame CA, USA) with 3,3’ -diaminobenzidine substrate and a haematoxylin counterstain. Staining was assessed in a semi-quantitative manner using a modified Quickscore, accounting for both extent and intensity of staining (Detre et al., 1995)

### Enzyme-linked immunosorbent assay (ELISA)

Secreted IL-6, IL-8 and IL-1β were quantified using the BD OptEIA(tm) (New Jersey, USA) sets for human IL-6, IL-8 and IL-1β detection. The procedure was performed following the manufacturer’s recommendation using 96-well plates. Absorbance was read at 450 nm within 30 minutes of adding the stop solution. Wavelength correction was done subtracting absorbance at 570 nm from absorbance at 450 nm.

### Quantitative (q) PCR

Total RNA was extracted from cell pellets using the Isolate II RNA Mini Kit (Bioline) RNA extraction Kit following manufacturer’s instructions. RNA was quantified using a Nanodrop 1000 Spectrophotometer (Thermo Fischer Scientific, Cambridge, UK). Five hundred nanograms of isolated RNA was reverse transcribed using the High Capacity cDNA Reverse Transcription Kit (Applied Biosystems) following manufacturer’s protocol using a Peltier Thermal cycler (MJ Research). cDNA was then stored at -20°C.

Gene expression was quantified with a Rotor-gene Q real-time PCR cycler (Qiagen, Manchester, UK) using SYBR green or TaqMan chemistry. Quantification was calculated using delta CT values normalized to either U6 or B2M. Each reaction was performed in triplicate. All reactions were performed in total volumes of 10 μl loading 500 ng of cDNA. The standard thermal cycle settings for a reaction consisted in 40 cycles including a melt curve analysis (when using SYBR green). One cycle consisted of: 95°C for 10 sec, 60°C for 15 sec, 72°C for 20 sec. SYBR green primers were designed in house and bought from Sigma-Aldrich (Gillingham, UK) corresponding to: (from 5’ to 3’, forward and reverse sequences) icIL1RA: CAGAAGACCTCCTGTCCTATGA, GAAGGTCTTCTGGTTAACATCCCAG, sIL-1RA: GAATGGAAATCTGCAGAGGCCTCCGC, GAAGGTCTTCTGGTTAACATCCCAG, IL-1R2: GTACGTGTTGGTAATGGGAGTTTC, CCGCTTGTAATGCCTCCCACGAAA, IFN-β: AAACTCATGAGCAGTCTGCA,, AGGAGATCTTCAGTTTCGGAGG, Lamin B1: CTCTCGTCGCATGCTGACAG, TCCCTTATTTCCGCCATCTCT. TaqMan probes were bought from Thermo Fischer Scientific (Cambridge, UK) and corresponded to: IL1RN (Hs: 00277299), IL-1R1 (Hs: 00991010), IL-1α (Hs: 00174092), IL-6 (Hs: 00985639), IL-8 (Hs: 00174103), B2M (Hs: 4325797).

### Western blotting

Protein was extracted by dissolving the cell pellets on an appropriate volume of lysis buffer on ice. Lysis buffer which consisted in one tablet of complete mini-EDTA free protease inhibitor cocktail (Roche, Basel, Switzerland) and one tablet of phosphatase inhibitors (PhosSTOP, Roche, Basel, Switzerland) dissolved in 10 ml RIPA Buffer (Sigma-Aldrich, Gillingham, UK). Cell suspensions were left for 30 minutes on ice and centrifuged at 13.300 rpm for 10 minutes at 4 °C. The supernatants were stored at -20 °C and the pellets were discarded. Protein quantification was done using the bicinchoninic acid assay (BCA) (Thermo Fischer Scientific, Cambridge, UK) according to the manufacturer’s protocol using a TECAN spectrophotometer (Spark).

Ten to fifty micrograms of protein were mixed with 2x SDS lysis buffer, heated for 5 minutes at 95°C and then loaded into 12-15% SDS-PAGE gels. Gels were run for ≈ 90 minutes at 150 V and transferred into nitrocellulose or PVDF membranes using the Trans-Blot® Turbo(tm) Transfer System (Bio-Rad, Deeside, UK) according to the manufacturer’s instructions. After transfer, membranes were blocked with 5% milk (Marvel) in TBS-T (Tris buffered saline 10mM, containing 0.5% Tween (v/v)) for 1 hour at room temperature in a rocking surface. The membranes were then incubated for 1 hour at room temperature or overnight at 4°C on a rocking platform with the primary antibody at the working concentration in 5% TBS-T milk. After incubation, membranes where washed three times for 10 minutes intervals with TBS-T and incubated with the secondary antibody at working concentration in 5% TBS-T milk for 1 hour at room temperature in a rocking platform. The membranes were then washed two times at intervals of 10 minutes with TBS-T and one time with TBS for 10 minutes and were ready for development. Membranes were developed with enhanced chemiluminescence (ECL), using Pierce ECL western blotting substrate (Thermo Fischer Scientific, Cambridge, UK), according to the manufacturer’s instructions. Signal was detected by exposure to an X-ray film (Thermo Fischer Scientific, Cambridge, UK) in a dark room and developed using a Compact X4 Developer (Xograph Imaging Systems) or alternatively using a Li-Cor C-Digit Western Blot Scanner and Image Studio Software. Densitometry was performed using the Image Studio Software. Bands were manually selected using the drawing tool and the intensity of each band was expressed normalized to the corresponding β-actin intensity.

Primary antibodies used were as follows: anti-IL-1RA (1: 2000, AF-280-NA, R&D Systems, Minneapolis, USA), anti-p16^Ink4a^ (1: 1000, 108349, Abcam, Cambridge, UK), anti-p21^Waf1/Cip1^ (1: 1000, MAB1047, R&D Systems, Minneapolis, USA), anti-phosphorylated p65 (1: 1000, 3033, Cell Signalling, London, UK), anti-IL-1α (1: 1000, Ab134908, Abcam, Cambridge, UK), anti-β-actin (1: 10000, A1978, Sigma-Aldrich, Gillingham, UK).

Secondary antibodies used were as follows: anti-mouse IgG HRP-conjugated (1: 5000, GTX221667-01, Genetex, California, USA), anti-goat IgG HRP-conjugated (1: 1000, HAF 017, R&D Systems, Minneapolis, USA), anti-rabbit IgG HRP-conjugated (1: 3000, 7074S, Cell signalling, London, UK).

### Statistical analysis

Statistical analysis was done using Graphpad Prism 7 Software. Comparison of two groups was done using the un-paired t-test. When the comparison included more than two groups, one-way ANOVA (analysis of variance) was performed. Two-way ANOVA was used when comparing two variables in multiple groups. A P-value < 0.05 was considered as statistically significant. The number of biological repeats is expressed as “N=” and the number of technical repeats as “n=”.

## ACKNOWLEDGMENTS

The authors would like to thank Prof Sheila Francis for advice and IL1RA reagents and Mrs Hayley Stanhope for preparing and cutting the histological sections

## AUTHOR CONTRIBUTION

K. D. Hunter and D. W. Lambert conceived and designed the project. S. Niklander, H. Crane and L. Darda performed the laboratory work. S. Niklander analysed the results. S. Niklander, K. D. Hunter and D. W. Lambert were involved in writing and all authors approved the final version of the manuscript.

## EXPANDED VIEW FIGURE LEGENDS

**Figure EV1:**
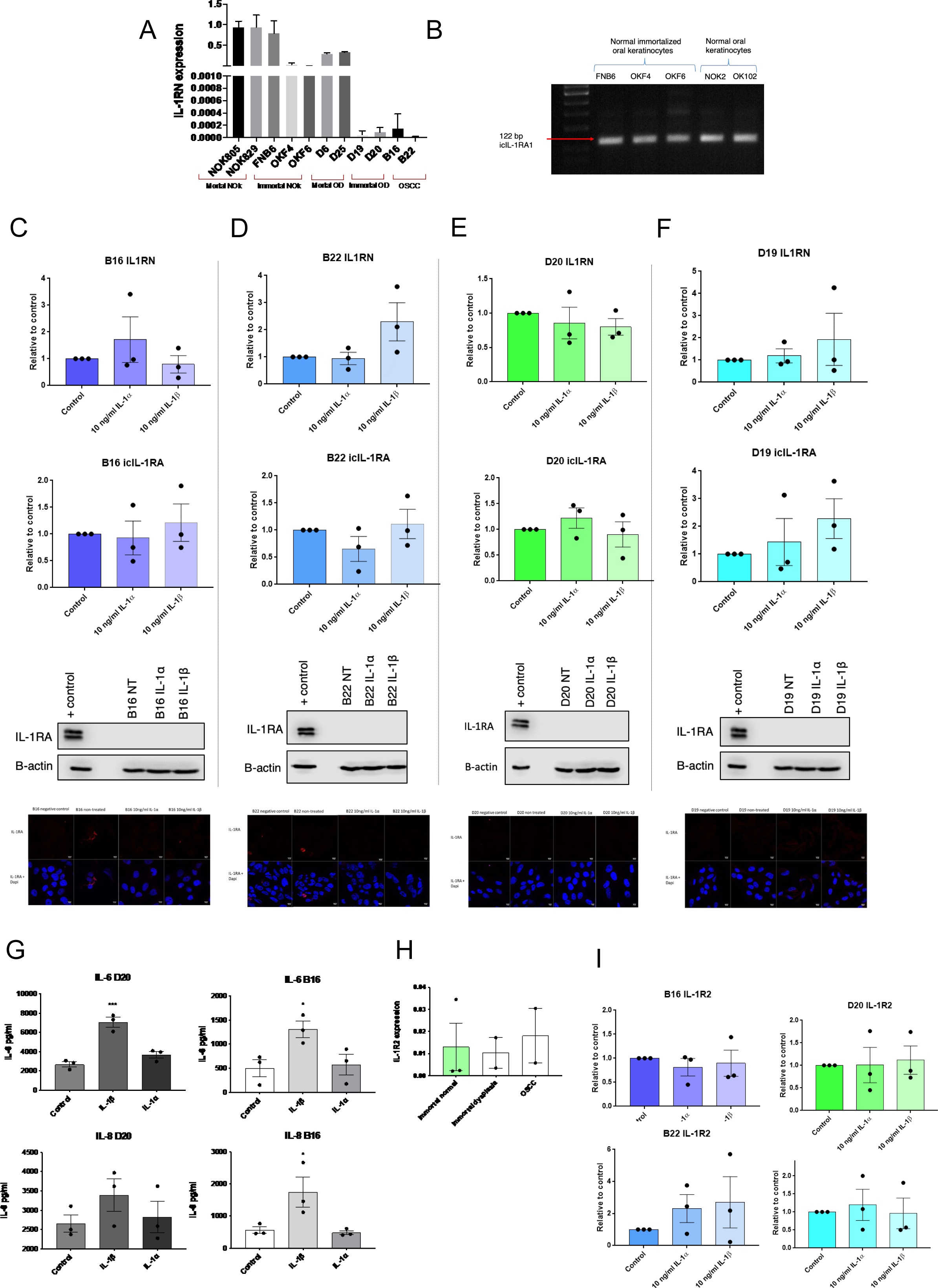
A: Expression of IL1RN mRNA levels per individual cell culture. B: PCR showing IL-1RA variants expressed in NOKs cell cultures. C-F IL1RN and icIL-1RA transcript levels and tIL-1RA expression after stimulation with 10 ng/ml of rIL-1α and 10 ng/ml of rIL-1β in OSCC (C, D) and OD (E, F) cell lines. G: IL-6 and IL-8 response after exposure to rIL-1α and rIL-1β in same samples from C-F. H: IL-1R2 mRNA expression in immortal NOKs (FNB6, OKF4 and OKF6), immortal OD (D19 and D20) and OSCC (b16 and B22) cell clines. Expression is express as fold change relative to the reference gene. Data are shown as mean ± SEM (N = 3 independent experiments, n = 3 technical replicates). One-way ANOVA with multiple comparisons was used to calculate the exact P value. I: IL-1R2 transcript levels after exposure to rIL-1α and rIL-1β in same samples from C-Data information: Data are shown as mean ± SEM (N = 3 independent experiments, n = 3 technical repeats).

**Figure EV2:**
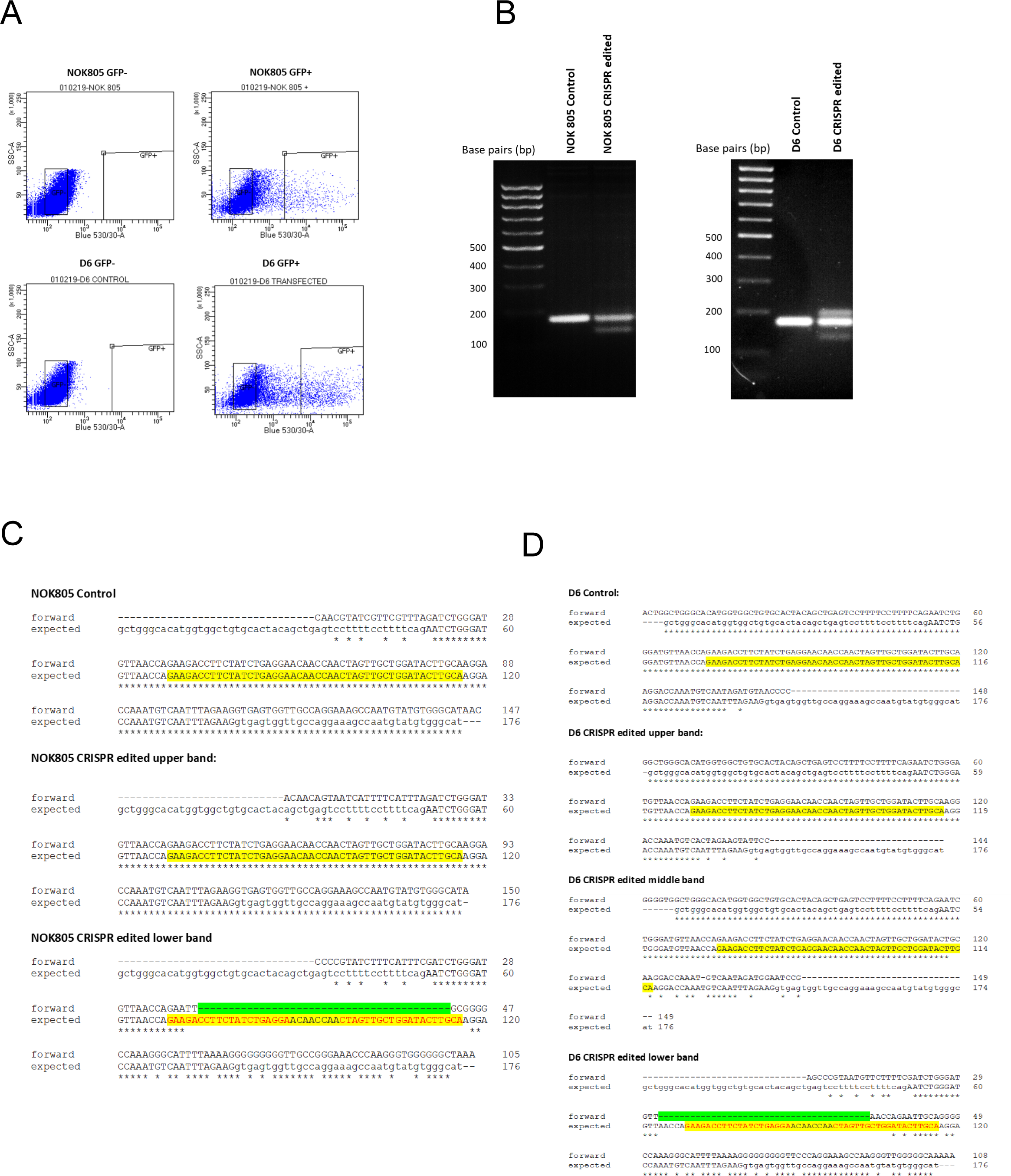
A: Transfected NOK805 and D6 cells were sorted into GFP+ and GFP-populations using a cell sorter. Same amount of GFP+ and GFP-cells per cell type were collected and expanded for further analysis. B: PCR products of NOK805 and D6 CRISPR edited and control cells using primers targeting the CRISPR target, resolved on a 2% agarose gel. Expected band size was of L 176 bp for non-edited cells and of L 130 bp for CRISPR edited cells. The presence of both bands in the CRISPR edited cells suggests an heterozygous population. C-D: Sequencing results of individual PCR products obtained after agarose electrophoresis of NOK805 and D6 edited cells. Expected deletions are highlighted in yellow. Acquired deletions are highlighted in green. Red letters correspond to each sgRNA target site. * means perfect match.

**Figure EV3:**
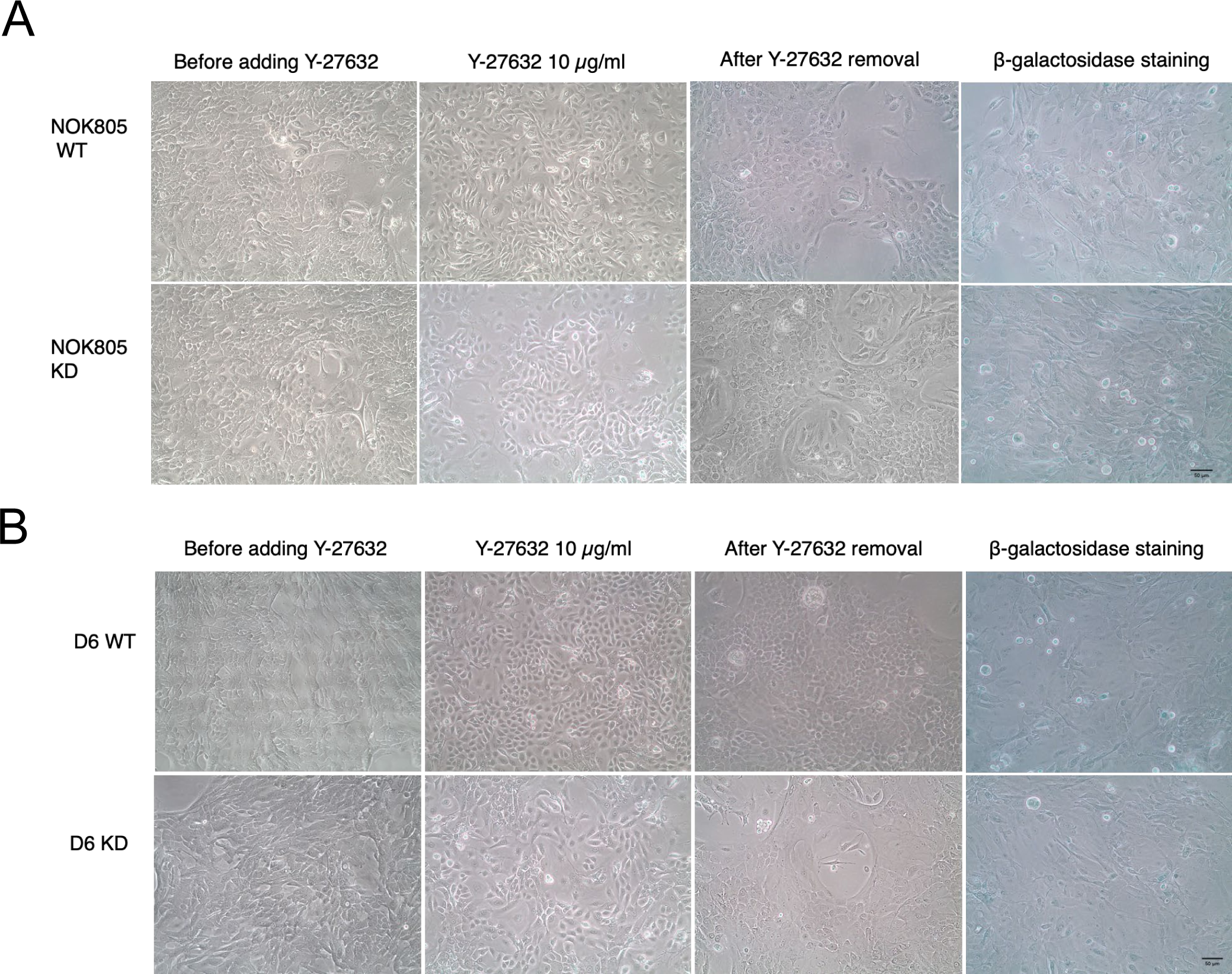
A-B: Shows cells morphology before and after adding Y-27632 to NOK805 (A) and D6 (B) cell cultures and SA-β-GAL activity after withdrawal.

**Figure EV4:**
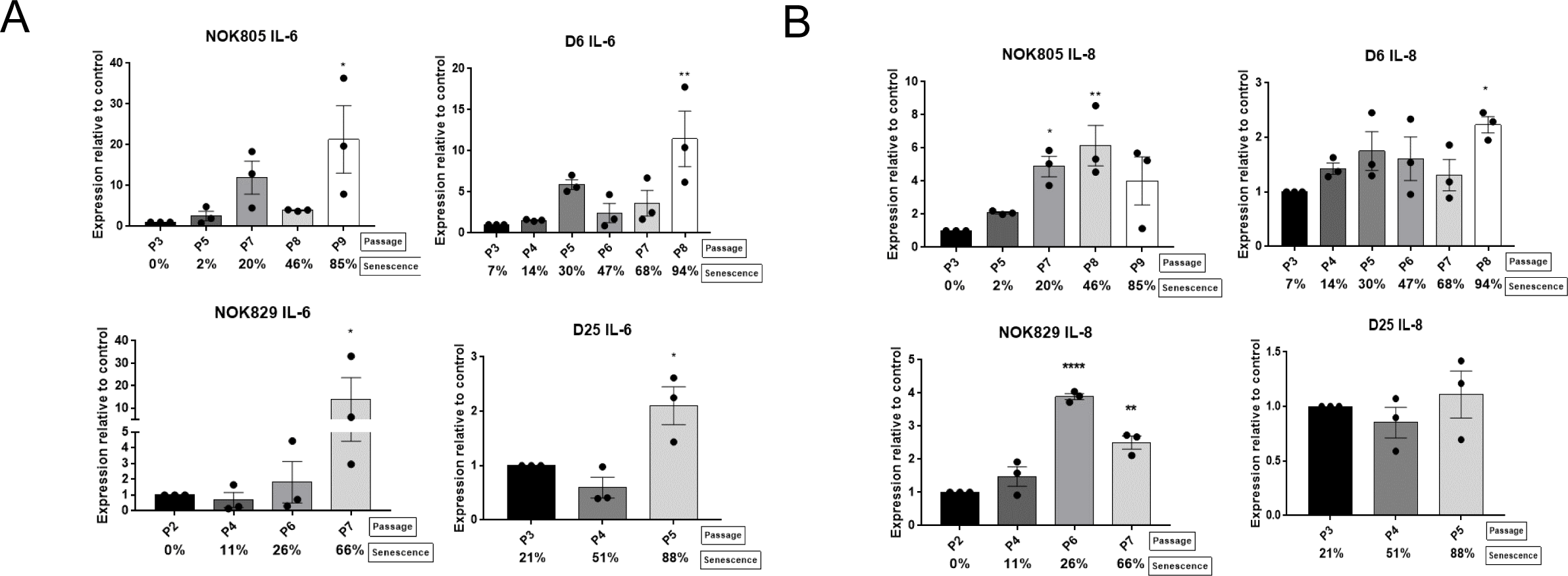
A-B: IL-6 and IL-8 mRNA transcript levels of normal and dysplastic oral keratinocytes during replicative senescence. Data are shown as mean fold change relative to control ± SEM (N = 3 independent experiments, n = 3 technical repeats). Statistical analysis was done using one-way ANOVA with multiple comparisons. Data information: *\*P < 0*.*05, **P < 0*.*005, *** P < 0*.*0005, **** P < 0*.*00001*

## REFERENCES

Abe, T., Maruyama, S., Yamazaki, M., Xu, B., Babkair, H., Sumita, Y., Cheng, J., Yamamoto, T. & Saku, T. 2017. Proteomic and histopathological characterization of the interface between oral squamous cell carcinoma invasion fronts and non-cancerous epithelia. Exp Mol Pathol, 102, 327–336.

Acosta, J. C., Banito, A., Wuestefeld, T., Georgilis, A., Janich, P., Morton, J. P., Athineos, D., Kang, T. W., Lasitschka, F., Andrulis, M., Pascual, G., Morris, K. J., Khan, S., Jin, H., Dharmalingam, G., Snijders, A. P., Carroll, T., Capper, D., Pritchard, C., Inman, G. J., Longerich, T., Sansom, O. J., Benitah, S. A., Zender, L. & Gil, J. 2013. A complex secretory program orchestrated by the inflammasome controls paracrine senescence. Nat Cell Biol, 15, 978–90.

Alevizos, I., Mahadevappa, M., Zhang, X., Ohyama, H., Kohno, Y., Posner, M., Gallagher, G. T., Varvares, M., Cohen, D., Kim, D., Kent, R., Donoff, R. B., Todd, R., Yung, C. M., Warrington, J. A. & Wong, D. T. 2001. Oral cancer in vivo gene expression profiling assisted by laser capture microdissection and microarray analysis. Oncogene, 20, 6196–204.

Ancrile, B., Lim, K. H. & Counter, C. M. 2007. Oncogenic Ras-induced secretion of IL6 is required for tumorigenesis. Genes Dev, 21, 1714–9.

Arend, W. R. 2002. The balance between IL-1 and IL-1Ra in disease. Cytokine & Growth Factor Reviews, 13, 323–340.

Campisi, J. 2005. Senescent cells, tumor suppression, and organismal aging: good citizens, bad neighbors. Cell, 120, 513–22.

Chapman, S., Mcdermott, D. H., Shen, K., Jang, M. K. & Mcbride, A. A. 2014. The effect of Rho kinase inhibition on long-term keratinocyte proliferation is rapid and conditional. Stem Cell Res Ther, 5, 60.

Cheng, W., Shivshankar, P., Zhong, Y., Chen, D., Li, Z. & Zhong, G. 2008. Intracellular interleukin-1alpha mediates interleukin-8 production induced by Chlamydia trachomatis infection via a mechanism independent of type I interleukin-1 receptor. Infect Immun, 76, 942–51.

Choi, P. & Chen, C. 2005. Genetic expression profiles and biologic pathway alterations in head and neck squamous cell carcinoma. Cancer, 104, 1113–28.

Colotta, F., Dower, S. K., Sims, J. E. & Mantovani, A. 1994. The type II ‘decoy’ receptor: a novel regulatory pathway for interleukin 1. Immunol Today, 15, 562–6.

Coppe, J. P., Patil, C. K., Rodier, F., Sun, Y., Munoz, D. P., Goldstein, J., Nelson, P. S., Desprez, P. Y. & Campisi, J. 2008. Senescence-associated secretory phenotypes reveal cell-nonautonomous functions of oncogenic RAS and the p53 tumor suppressor. PLoS Biol, 6, 2853–68.

Cromer, A., Carles, A., Millon, R., Ganguli, G., Chalmel, F., Lemaire, F., Young, J., Dembele, D., Thibault, C., Muller, D., Poch, O., Abecassis, J. & Wasylyk, B. 2004. Identification of genes associated with tumorigenesis and metastatic potential of hypopharyngeal cancer by microarray analysis. Oncogene, 23, 2484–98.

Davalos, A. R., Coppe, J. P., Campisi, J. & Desprez, P. Y. 2010. Senescent cells as a source of inflammatory factors for tumor progression. Cancer Metastasis Rev, 29, 273–83.

Detre, S., Saclani Jotti, G. & Dowsett, M. 1995. A “quickscore” method for immunohistochemical semiquantitation: validation for oestrogen receptor in breast carcinomas. J Clin Pathol, 48, 876–8.

Dinarello, C. A. 2010. Why not treat human cancer with interleukin-1 blockade? Cancer Metastasis Rev, 29, 317–29.

Dou, Z., Ghosh, K., Vizioli, M. G., Zhu, J., Sen, P., Wangensteen, K. J., Simithy, J., Lan, Y., Lin, Y., Zhou, Z., Capell, B. C., Xu, C., Xu, M., Kieckhaefer, J. E., Jiang, T., Shoshkes-Carmel, M., Tanim, K., Barber, G. N., Seykora, J. T., Millar, S. E., Kaestner, K. H., Garcia, B. A., Adams, P. D. & Berger, S. L. 2017. Cytoplasmic chromatin triggers inflammation in senescence and cancer. Nature, 550, 402–406.

Duffey, D. C., Chen, Z., Dong, G., Ondrey, F. G., Wolf, J. S., Brown, K., Siebenlist, U. & Van Waes, C. 1999. Expression of a dominant-negative mutant inhibitor-kappaBalpha of nuclear factor-kappaB in human head and neck squamous cell carcinoma inhibits survival, proinflammatory cytokine expression, and tumor growth in vivo. Cancer Res, 59, 3468–74.

Eisenberg, S. P., Evans, R. J., Arend, W. P., Verderber, E., Brewer, M. T., Hannum, C. H. & Thompson, R. C. 1990. Primary structure and functional expression from complementary DNA of a human interleukin-1 receptor antagonist. Nature, 343, 341–6.

Garat, C. & Arend, W. P. 2003. Intracellular IL-1Ra type 1 inhibits IL-1-induced IL-6 and IL-8 production in Caco-2 intestinal epithelial cells through inhibition of p38 mitogen-activated protein kinase and NF-kappaB pathways. Cytokine, 23, 31–40.

Gluck, S., Guey, B., Gulen, M. F., Wolter, K., Kang, T. W., Schmacke, N. A., Bridgeman, A., Rehwinkel, J., Zender, L. & Ablasser, A. 2017. Innate immune sensing of cytosolic chromatin fragments through cGAS promotes senescence. Nat Cell Biol, 19, 1061–1070.

Goertzen, C., Mahdi, H., Laliberte, C., Meirson, T., Eymael, D., Gil-Henn, H. & Magalhaes, M. 2018. Oral inflammation promotes oral squamous cell carcinoma invasion. Oncotarget, 9, 29047–29063.

Guo, Y., Ayers, J. L., Carter, K. T., Wang, T., Maden, S. K., Edmond, D., Newcomb, P. P., Li, C., Ulrich, C., Yu, M. & Grady, W. M. 2019. Senescence-associated tissue microenvironment promotes colon cancer formation through the secretory factor GDF15. Aging Cell, 18, e13013.

Hakelius, M., Reyhani, V., Rubin, K., Gerdin, B. & Nowinski, D. 2016. Normal Oral Keratinocytes and Head and Neck Squamous Carcinoma Cells Induce an Innate Response of Fibroblasts. Anticancer Res, 36, 2131–7.

Hannum, C. H., Wilcox, C. J., Arend, W. P., Joslin, F. G., Dripps, D. J., Heimdal, P. L., Armes, L. G., Sommer, A., Eisenberg, S. P. & Thompson, R. C. 1990. Interleukin-1 receptor antagonist activity of a human interleukin-1 inhibitor. Nature, 343, 336–343.

Haskill, S., Martin, G., Van Le, L., Morris, J., Peace, A., Bigler, C. F., Jaffe, G. J., Hammerberg, C., Sporn, S. A., Fong, S. & Arend, W. P. 1991. cDNA cloning of an intracellular form of the human interleukin 1 receptor antagonist associated with epithelium. Cell Biology, 88, 3681–3685.

Hunter, K. D., Thurlow, J. K., Fleming, J., Drake, P. J., Vass, J. K., Kalna, G., Higham, D. J., Herzyk, P., Macdonald, D. G., Parkinson, E. K. & Harrison, P. R. 2006. Divergent routes to oral cancer. Cancer Res, 66, 7405–13.

Jang Da, H., Bhawal, U. K., Min, H. K., Kang, H. K., Abiko, Y. & Min, B. M. 2015. A transcriptional roadmap to the senescence and differentiation of human oral keratinocytes. J Gerontol A Biol Sci Med Sci, 70, 20–32.

Koike, H., Uzawa, K., Nakashima, D., Shimada, K., Kato, Y., Higo, M., Kouzu, Y., Endo, Y., Kasamatsu, A. & Tanzawa, H. 2005. Identification of differentially expressed proteins in oral squamous cell carcinoma using a global proteomic approach. Int J Oncol, 27, 59–67.

Kuilman, T., Michaloglou, C., Vredeveld, L. C., Douma, S., Van Doorn, R., Desmet, C. J., Aarden, L. A., Mooi, W. J. & Peeper, D. S. 2008. Oncogene-induced senescence relayed by an interleukin-dependent inflammatory network. Cell, 133, 1019–31.

Kurzrock, R., Hickish, T., Wyrwicz, L., Saunders, M., Wu, Q., Stecher, M., Mohanty, P., Dinarello, C. A. & Simard, J. 2019. Interleukin-1 receptor antagonist levels predict favorable outcome after bermekimab, a first-in-class true human interleukin-1alpha antibody, in a phase III randomized study of advanced colorectal cancer. Oncoimmunology, 8, 1551651.

La, E., Rundhaug, J. E. & Fischer, S. M. 2001. Role of intracellular interleukin-1 receptor antagonist in skin carcinogenesis. Mol Carcinog, 30, 218–23.

Lallemant, B., Evrard, A., Combescure, C., Chapuis, H., Chambon, G., Raynal, C., Reynaud, C., Sabra, O., Joubert, D., Hollande, F., Lallemant, J. G. LUMBROSO, & Brouillet, J. P. 2009. Clinical relevance of nine transcriptional molecular markers for the diagnosis of head and neck squamous cell carcinoma in tissue and saliva rinse. BMC Cancer, 9, 370.

Lau, L., Porciuncula, A., Yu, A., Iwakura, Y. & David, G. 2019. Uncoupling the Senescence-Associated Secretory Phenotype from Cell Cycle Exit via Interleukin-1 Inactivation Unveils Its Protumorigenic Role. Mol Cell Biol, 39.

Leethanakul, C., Patel, V., Gillespie, J., Shillitoe, E., Kellman, R. M., Ensley, J. F., Limwongse, V., Emmert-Buck, M. R., Krizman, D. B. & Gutkind, J. S. 2000. Gene expression profiles in squamous cell carcinomas of the oral cavity: use of laser capture microdissection for the construction and analysis of stage-specific cDNA libraries. Oral Oncol, 36, 474–83.

Leung, E. Y., Mcmahon, J. D., Mclellan, D. R., Syyed, N., Mccarthy, C. E., Nixon, C., Orange, C., Brock, C., Hunter, K. D. & Adams, P. D. 2017. DNA damage marker phosphorylated histone H2AX is a potential predictive marker for progression of epithelial dysplasia of the oral cavity. Histopathology, 71, 522–528.

Lewis, A. M., Varghese, S., Xu, H. & Alexander, H. R. 2006. Interleukin-1 and cancer progression: the emerging role of interleukin-1 receptor antagonist as a novel therapeutic agent in cancer treatment. J Transl Med, 4, 48.

Loaiza, N. & Demaria, M. 2016. Cellular senescence and tumor promotion: Is aging the key? Biochim Biophys Acta, 1865, 155–167.

Lodi, G., Franchini, R., Warnakulasuriya, S., Varoni, E. M., Sardella, A., Kerr, A. R., Carrassi, A., Macdonald, L. C. & Worthington, H. V. 2016. Interventions for treating oral leukoplakia to prevent oral cancer. Cochrane Database Syst Rev, 7, Cd001829.

Malaquin, N., Vercamer, C., Bouali, F., Martien, S., Deruy, E., Wernert, N., Chwastyniak, M., Pinet, F., Abbadie, C. & Pourtier, A. 2013. Senescent fibroblasts enhance early skin carcinogenic events via a paracrine MMP-PAR-1 axis. PLoS One, 8, e63607.

Malyak, M., Guthridge, J. M., Hance, K. R., Dower, S. K., Freed, J. H. & Arend, W. P. 1998a. Characterization of a low molecular weight isoform of IL-1 receptor antagonist. J Immunol, 161, 1997–2003.

Malyak, M., Smith, M. F., jr., Abel, A. A., Hance, K. R. & Arend, W. P. 1998b. The differential production of three forms of IL-1 receptor antagonist by human neutrophils and monocytes. J Immunol, 161, 2004–10.

Marur, S. & Forastiere, A. A. 2016. Head and Neck Squamous Cell Carcinoma: Update on Epidemiology, Diagnosis, and Treatment. Mayo Clin Proc, 91, 386–96.

Mcgregor, F., Muntoni, A., Fleming, J., Brown, J., Felix, D. H., Macdonald, D. G., Parkinson, E. K. & Harrison, P. R. 2002. Molecular changes associated with oral dysplasia progression and acquisition of immortality: potential for its reversal by 5-azacytidine. Cancer Res, 62, 4757–66.

Mcgregor, F., Wagner, E., Felix, D., Soutar, D., Parkinson, K. & Harrison, P. R. 1997. Inappropriate retinoic acid receptor-beta expression in oral dysplasias: correlation with acquisition of the immortal phenotype. Cancer Res, 57, 3886–9.

Merhi-Soussi, F., Berti, M., Wehrle-Haller, B. & Gabay, C. 2005. Intracellular interleukin-1 receptor antagonist type 1 antagonizes the stimulatory effect of interleukin-1 alpha precursor on cell motility. Cytokine, 32, 163–70.

Napier, S. S. & Speight, P. M. 2008. Natural history of potentially malignant oral lesions and conditions: an overview of the literature. J Oral Pathol Med, 37, 1–10.

Natarajan, E., Omobono, J. D., 2nd, Guo, Z., Hopkinson, S., Lazar, A. J., Brenn, T., Jones, J. C. & Rheinwald, J. G. 2006. A keratinocyte hypermotility/growth-arrest response involving laminin 5 and p16INK4A activated in wound healing and senescence. Am J Pathol, 168, 1821–37.

Natarajan, E., Saeb, M., Crum, C. P., Woo, S. B., Mckee, P. H. & Rheinwald, J. G. 2003. Co-expression of p16(INK4A) and laminin 5 gamma2 by microinvasive and superficial squamous cell carcinomas in vivo and by migrating wound and senescent keratinocytes in culture. Am J Pathol, 163, 477–91.

Orjalo, A. V., Bhaumik, D., Gengler, B. K., Scott, G. K. & Campisi, J. 2009. Cell surface-bound IL-1alpha is an upstream regulator of the senescence-associated IL-6/IL-8 cytokine network. Proc Natl Acad Sci U S A, 106, 17031–6.

Ortiz-Montero, P., Londono-Vallejo, A. & Vernot, J. P. 2017. Senescence-associated IL-6 and IL-8 cytokines induce a self- and cross-reinforced senescence/inflammatory milieu strengthening tumorigenic capabilities in the MCF-7 breast cancer cell line. Cell Commun Signal, 15, 17.

Palmer, G., Trolliet, S., Talabot-Ayer, D., Mezin, F., Magne, D. & Gabay, C. 2005. Pre-interleukin-1alpha expression reduces cell growth and increases interleukin-6 production in SaOS-2 osteosarcoma cells: Differential inhibitory effect of interleukin-1 receptor antagonist (icIL-1Ra1). Cytokine, 31, 153–60.

Rheinwald, J. G. & Green, H. 1975. Serial cultivation of strains of human epidermal keratinocytes: the formation of keratinizing colonies from single cells. Cell, 6, 331–43.

Rheinwald, J. G., Hahn, W. C., Ramsey, M. R., Wu, J. Y., Guo, Z., Tsao, H., De Luca, M., Catricala, C. & O’Toole, K. M. 2002. A Two-Stage, p16INK4A- and p53-Dependent Keratinocyte Senescence Mechanism That Limits Replicative Potential Independent of Telomere Status. Molecular and Cellular Biology, 22, 5157–5172.

Rovillain, E., Mansfield, L., Caetano, C., Alvarez-Fernandez, M., Caballero, O. L., Medema, R. H., Hummerich, H. & Jat, P. S. 2011. Activation of nuclear factor-kappa B signalling promotes cellular senescence. Oncogene, 30, 2356–66.

Schmalbach, C. E., Chepeha, D. B., Giordano, T. J., Rubin, M. A., Teknos, T. N., Bradford, C. R., Wolf, G. T., Kuick, R., Misek, D. E., Trask, D. K. & Hanash, S. 2004. Molecular profiling and the identification of genes associated with metastatic oral cavity/pharynx squamous cell carcinoma. Arch Otolaryngol Head Neck Surg, 130, 295–302.

Shiiba, M., Saito, K., Yamagami, H., Nakashima, D., Higo, M., Kasamatsu, A., Sakamoto, Y., Ogawara, K., Uzawa, K., Takiguchi, Y. & Tanzawa, H. 2015. Interleukin-1 receptor antagonist (IL1RN) is associated with suppression of early carcinogenic events in human oral malignancies. Int J Oncol, 46, 1978–84.

Sparmann, A. & Bar-Sagi, D. 2004. Ras-induced interleukin-8 expression plays a critical role in tumor growth and angiogenesis. Cancer Cell, 6, 447–58.

Tomo, S., Biss, S. P., Crivelini, M. M., De Oliveira, S. H. P., Biasoli, E. R., Tjioe, K. C., Bernabe, D. G., Villa, L. L. & Miyahara, G. I. 2020. High p16(INK4a) immunoexpression is not HPV dependent in oral leukoplakia. Arch Oral Biol, 115, 104738.

Uekawa, N., Nishikimi, A., Isobe, K., Iwakura, Y. & Maruyama, M. 2004. Involvement of IL-1 family proteins in p38 linked cellular senescence of mouse embryonic fibroblasts. FEBS Lett, 575, 30–4.

Villa, A., Celentano, A., Glurich, I., Borgnakke, W. S., Jensen, S. B., Peterson, D. E., Delli, K., Ojeda, D., Vissink, A. & Farah, C. S. 2019. World Workshop on Oral Medicine VII: Prognostic biomarkers in oral leukoplakia: A systematic review of longitudinal studies. Oral Dis, 25 Suppl 1, 64–78.

Vincent, J., Adura, C., Gao, P., Luz, A., Lama, L., Asano, Y., Okamoto, R., Imaeda, T., Aida, J., Rothamel, K., Gogakos, T., Steinberg, J., Reasoner, S., Aso, K., Tuschl Patel, D. J., Glickman, J. F. & Ascano, M. 2017. Small molecule inhibition of cGAS reduces interferon expression in primary macrophages from autoimmune mice. Nat Commun, 8, 750.

Von Biberstein, S. E., Spiro, J. D., Lindquist, R. & Kreutzer, D. L. 1996. Interleukin-1 Receptor Antagonist in Head and Neck Squamous Cell Carcinoma. Arch Otolaryngol Head Neck Surg, 122, 751–759.

Weissbach, L., Tran, K., Colquhoun, S. A., Champliaud, M. F. & Towle, C. A. 1998. Detection of an interleukin-1 intracellular receptor antagonist mRNA variant. Biochem Biophys Res Commun, 244, 91–5.

Werman, A., Werman-Venkert, R., White, R., Lee, J. K., Werman, B., Krelin, Y., Voronov, E., Dinarello, C. A. & Apte, R. N. 2004. The precursor form of IL-1alpha is an intracrine proinflammatory activator of transcription. Proc Natl Acad Sci U S A, 101, 2434–9.

Whipple, M. E., Mendez, E., Farwell, D. G., Agoff, S. N. & Chen, C. 2004. A genomic predictor of oral squamous cell carcinoma. Laryngoscope, 114, 1346–54.

Wiggins, K. A., Parry, A. J., Cassidy, L. D., Humphry, M., Webster, S. J., Goodall, J. C., Narita, M. & Clarke, M. C. H. 2019. IL-1alpha cleavage by inflammatory caspases of the noncanonical inflammasome controls the senescence-associated secretory phenotype. Aging Cell, 18, e12946.

Wolf, J. S., Chen, Z., Dong, G., Sunwoo, J. B., Bancroft, C. C., Capo, D. E., Yeh, N. T., Mukaida, N. & Van Waes, C. 2001. IL (Interleukin)-1 Promotes Nuclear Factor-B and AP-1-induced IL-8 Expression, Cell Survival, and Proliferation in Head and Neck Squamous Cell Carcinomas. Clinical Cancer Research, 7, 1812–1820.

Wu, S., Hu, G., Chen, J. & Xie, G. 2014. Interleukin 1beta and interleukin 1 receptor antagonist gene polymorphisms and cervical cancer: a meta-analysis. Int J Gynecol Cancer, 24, 984–90.

Wu, T., Hong, Y., Jia, L., Wu, J., Xia, J., Wang, J., Hu, Q. & Cheng, B. 2016. Modulation of IL-1beta reprogrammes the tumor microenvironment to interrupt oral carcinogenesis. Sci Rep, 6, 20208.

Zhang, Y., Liu, C., Peng, H., Zhang, J. & Feng, Q. 2012. IL1 receptor antagonist gene IL1-RN variable number of tandem repeats polymorphism and cancer risk: a literature review and meta-analysis. PLoS One, 7, e46017.

